# A vestibulospinal pathway for context-dependent motor control of the mouse tail

**DOI:** 10.1101/2024.12.25.630321

**Authors:** Salvatore Andrea Lacava, Necmettin Isilak, Nishtha Ranawat, Julian Katzke, Johannes Frederik Hugo Hoedemaker, Yutaka Yoshida, Marylka Yoe Uusisaari

## Abstract

The tail movement is critical for maintaining balance during locomotion in many animal species, yet its underlying neuro-muscular control remains poorly understood. In this study we investigated what are the neuronal substrates responsible for tail control in mice. Using high-resolution microCT scans and retrograde labeling, we lay out the neuro-muscular organization of the tail and identified distinct pools of motoneurons in the spinal cord that innervate proximal and distal muscles. We further show that the spinal vestibular nucleus (SpVN) in the brainstem sends direct projections to the same spinal cord segments where tail motoneurons are located. The activation of these vestibulospinal neurons using optogenetics results in more accurate tail movements during challenging balance tasks. Our results reveal that the vestibular system’s influence on tail control is context-dependent, enhancing balance performance under uncertain sensory conditions. These findings provide novel insights into the neural circuits responsible for maintaining balance, highlighting the role of the vestibulospinal pathway in context-dependent modulation of tail movement to maintain stability during complex locomotor tasks.

## INTRODUCTION

Balance and context-appropriate posture are fundamental to all animal movement and behavior, and have been so throughout animal evolution. When experiencing unexpected and undesired changes in posture or balancing state, animals rapidly engage muscles appropriate to the external and internal context (^1,2^). While the repertoire of corrective movements available to an animal has expanded alongside increasing complexity of animals’ body plans and their living environments (e.g. evolution of upright legged posture, arboreal ecotypes;^3^), tail movements are likely among the most ancient and fundamental balancing responses throughout the chordate lineage (^4,5^).

Several recent studies have described how various animal species use their tails to maintain balance and posture in different contexts (^6,7,8^), and our previous work (^9^) demonstrated how laboratory mice mitigate roll-plane balance perturbations by rotating their tails and generating compensatory angular momentum. Notably, even though mice can and do engage other stabilizing movements when their balance is disturbed (^10^), the tail offers a more tractable model to understand balancing strategies due to its clear visibility (large tail swings compared to subtle activation of leg extensor musculature). Furthermore, mice use their tails primarily for balance, whereas their limbs serve multiple functions - supporting weight, enabling locomotion, and aiding in balance - making it challenging to separate balancing responses from other movements.

One limitation in leveraging the tail’s advantages in behavioral neuroscience is our limited understanding of the neuromuscular organization of the tail. Specifically, despite the relative structural simplicity of the mouse tail, very little description of the tail muscle architecture is available, and even the location of tail-controlling motoneurons at the level of spinal segments remains elusive. Without this knowledge it is not clear whether descending supra-spinal pathways - such as the vestibulo-spinal tract directly involved in other balancing responses (e.g. vestibulo-collic reflex,^11,12,13^) - drive tail movements, or if the tail is mainly controlled by spinal reflexes without the need for supraspinal input.

In this study we localize tail-controlling motoneurons (tail-MNs) in the sacral and coccygeal spinal segments (sacral (S)3, S4, coccygeal (Co)1-Co3) and establish the existence and function of a descending vestibular pathway that targets the tail-related spinal segments. This tail-targeting vestibular pathway originates predominantly from the Spinal Vestibular Nucleus (SpVN), and is distinct from the two previously-described vestibulospinal pathways originating from the medial and lateral vestibular nuclei (^14,15^).

To examine the behavioral significance of the sacral-vestibulospinal pathway, we expressed channelrhodopsin (ChR) in vestibular neurons and tracked tail movements evoked in response to optogenetic vestibular activation while crossing a narrow platform (hereafter referred to as the ridge task, previously described in^9^). Intriguingly, indiscriminate activation of vestibular neurons drove complex, whole-body responses that also involved tail movements, while selective activation of the sacral-projecting vestibulospinal neurons resulted in much subtler and tail-specific effects that displayed clear context-dependence. Finally, combining the selective optogenetic stimulation with sub-threshold balance perturbations in the roll-plane improved the selection of appropriate compensatory tail responses.

Our results call for attention not only to the the neuronal control of tail movements but also to the contribution of the vestibular system to fine-tune postural responses during locomotion.

## RESULTS

### Anatomy of mouse tail and distribution of tail-motoneurons

To our knowledge, detailed anatomical studies describing the mouse tail morphology have not been reported. Thus, to guide our investigation on neuronal circuits involved in tail motor control, we acquired a microCT-based reconstruction of the extrinsic-to-mid regions of the tail (12-week-old male mouse C57BL/6; Figure 1 A). The reconstruction highlights the multi-scale arrangement of muscles with respect to the vertebrae: the extrinsic muscles (located at the tail base) span over several vertebrae forming extrinsic units, while the distal muscles only reach over a single vertebral joint. This arrangement, known as metameric arrangement, resembles the forelimb organization of most mammals (^16^), and suggests that extrinsic musculature is capable of generating the force needed for large movements, while the intrinsic muscles are used for more fine movements (for example the coiling motion observed when mice wrap their tails around objects). Moreover, the long tendons packed between extrinsic muscles could store significant amounts of elastic energy, which is a function necessary to produce fast propulsive movements (such as the hindlimb tendons of kangaroo rats^17^).

**Figure 1:**
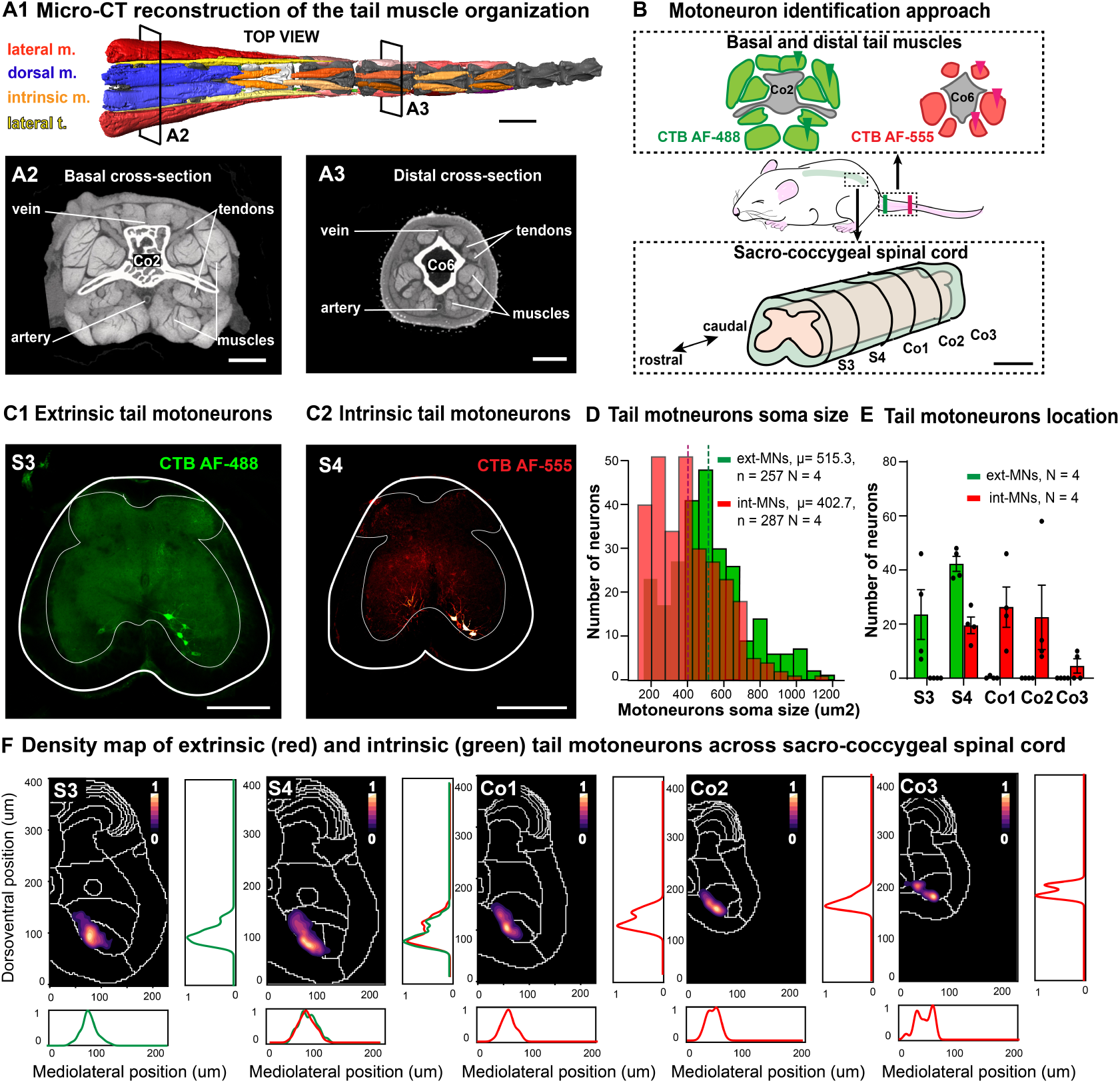
Tail-motoneurons targeting extrinsic and intrinsic muscles reside in different spinal segments. **A:** Micro-CT scan of the basal-to-mid mouse tail reveals multiscale organization of the musculature and connective elements. **A1:** 3D reconstruction of the tail (top view, from coccygeal vertebra Co2 to Co10) highlighting the metameric arrangement of extrinsic and intrinsic musculature. Red and blue, proximal (extrinsic) muscles; orange, distal (intrinsic) muscles. The lateral tendons of extrinsic muscles are displayed in yellow. Note that for visualizing the vertebrae, the distal most muscles are omitted from the graphic. **A2-A3:** single cross-section images from the micro-CT scan from locations indicated in A1. **B:** Schematic depiction of the approach for revealing tail-motoneuron location. Note that spinal and vertebral nomenclature do not align in caudal parts (e.g. spinal segment Co2 does not reside within vertebra Co2;^49^). **C:** Example of retrograde labeling revealed with injection to extrinsic (C1) and intrinsic (C2) tail muscles, shown as standard deviation projection images. **D:** Intrinsic-projecting motoneurons include smaller soma sizes than extrinsic-projecting motoneurons. **E:** Rostrocaudal distribution of tail-motoneuron somata in the spinal cord correlates with extrinsic-intrinsic location of the muscles. Each dot represents data from a single animal (N=4). **F:** Density maps of tail-motoneuron locations from 4 animals. Each map shows density of pooled extrinsic and intrinsic-projecting tail-motoneurons; the individual spatial distributions of extrinsic and intrinsic populations are shown with green and red histograms outside the maps. Scale bars; A1, 5mm; A2-A3, 1mm; B, 500 µm; C1-C2, 200 µm.

Next, to localize the motoneurons controlling tail muscles, we injected green and red retrograde tracers to sets of extrinsic and intrinsic muscles of four 10-week-old male mice, (respectively CTB AF-488 and CTB AF-555; Figure 1 B). The tissue was collected 96 hours later, and the green or red-labeled neurons were revealed in the ventral laminae of spinal cord segments S3-Co3 (Figure 1 C). Intriguingly, intrinsic-tail-motoneurons (Tail-MN_int_) had significantly smaller somata than the extrinsic-tail motoneurons (Tail-MN_ext_) (Tail-MN_int_ = 402.7 +-10.69 vs Tail-MN_ext_ = 515.3+-13.02, unpaired t-test p *<* 0.001), and they reside in more caudal segments of the spinal cord with little overlap only in S4 (Figure 1 D-E). Despite these differences, both proximal and distal-targeting tail-motoneurons were found in nearly-identical medial region of the spinal cord laminae VIII and IX ( Figure 1 F). No labeled motoneurons were ever seen more rostral than S3.

### Distribution of vestibulospinal axons in the sacral spinal cord

To examine whether the vestibular complex neurons project to the sacral spinal cord segments where the tail-MN_ext_ were found, we used a retrograde intersectional viral labeling approach (see Methods, and Figure 2 A). As shown in the examples in Figure 2 A1-A3, the axons could be clearly identified, allowing the reconstruction of density maps (Figure 2 B1) within the same co-ordinate system as was used in Figure 1 B to map tail-motoneuron locations. The most densely labeled region extended from lamina VII up to IX (quantified in Figure 2 B2) highlighting that the vestibular projection largely terminates among the interneuronal circuits of the spinal cord. However, combining the intersectional retrograde labeling of vestibular axons with retrograde labeling from the tail muscle (Figure 2 C) showed that the tail-motoneurons do also receive direct contacts from the vestibular neurons, and the existence of a direct connection between vestibular nuclei and tail-MNs was confirmed with transynaptic tracing (Suppl. Figure 2). Specifically, we injected in the tail N2cdelG-tdTomato rabies virus, as well as AAV6-oG-P2A-H2BStayGold (see Methods), which labeled tail-MNs as expected, as well as interneurons in sacral and lumbar spinal cord, and neurons in the lateral vestibular nucleus (Supp. Figure 2). We also noted a high density of the vestibulo-spinal axons within the spinal precerebellar nucleus (Figure 2 B2).

**Figure 2:**
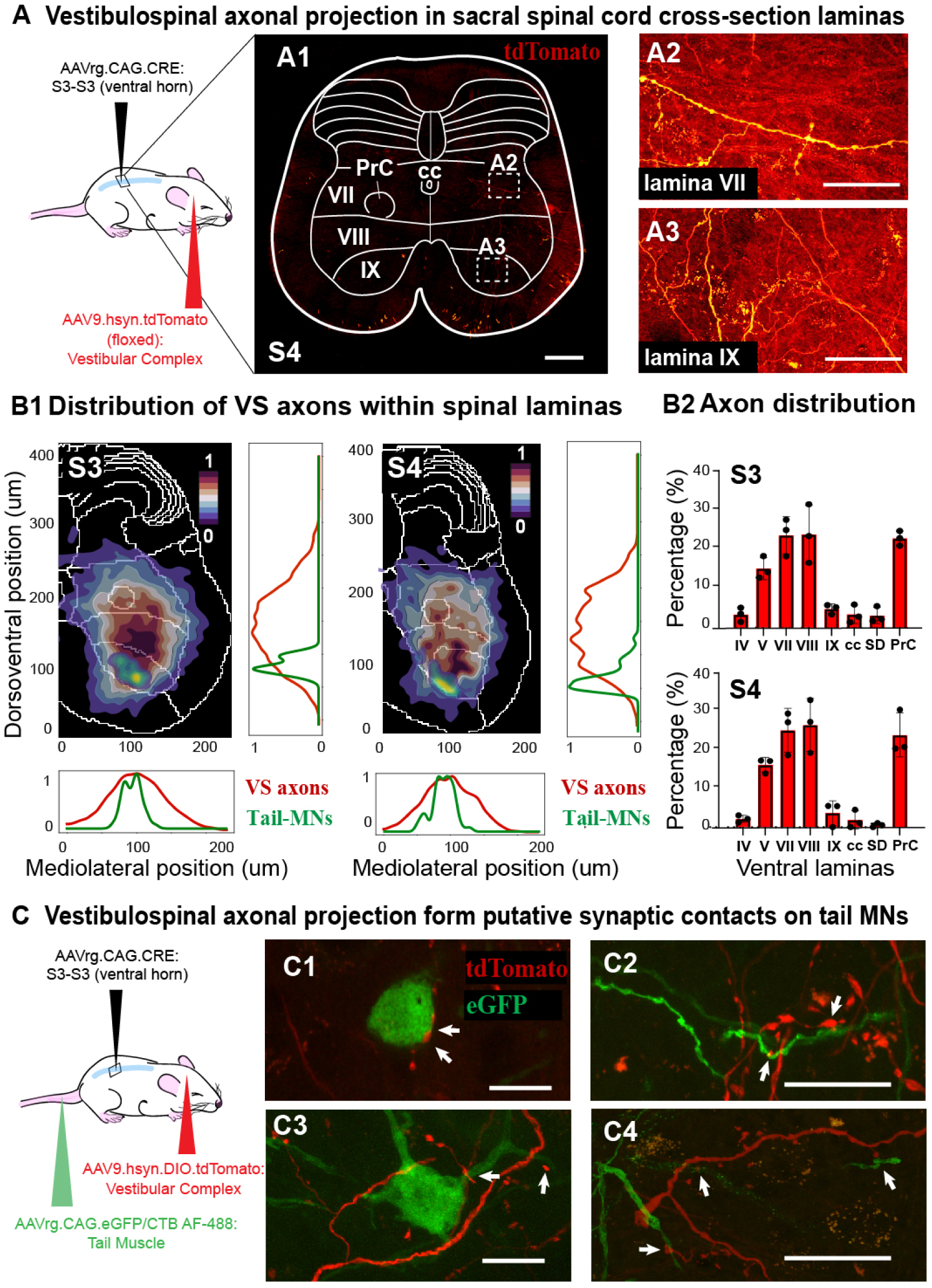
Tail-related vestibular neurons project to the intermediate lamina of the spinal cord but also contact tail-motoneurons directly. **A1:** Schematic of the intersectional viral approach. **A2:** A confocal image of S3 spinal cord section with tdTomato-labeled axons, overlaid with schematic definition of laminae. The rectangles indicate positions of higher-magnified images shown in A3 and A4. **B:** Anatomical data from 3 animals compiled into axonal density maps (B1), combined with the tail-motoneuron density information as shown in Figure 1B. Graphs on the side and bottom of maps display localization in dorsoventral and mediolateral planes, respectively. **B2:** Summary of axonal density distribution among the ventral lamina (IV-IX). **C:** Image of putative synaptic contacts between vestibular neurons and tail-motoneurons. **C1:** Schematic of the viral strategy; C2-4, examples of motoneuron somata (left) and dendrites (right) found at close apposition to vestibulospinal axons. The images are from lamina IX. Scale bars: A1-A3: 100 µm; C1-C4: 20 µm. Abbreviations: cc, central canal; PrC, pre-cerebellar nucleus; VII, lamina seven; VIII, lamina eight; IX, lamina nine.

### Distribution of vestibulospinal neurons projecting to sacral spinal cord

After having confirmed the existence of vestibulospinal neurons targeting the sacral segments of the spinal cord, we further examined the distribution of their soma within the vestibular complex. As shown in Figure 3A, cre-dependent expression of tdTomato was clearly visible in numerous vestibular neurons when paired with retrogradely-transported cre-expression virus. Combining samples from 3 animals with this labeling procedure and registering the data to the Allen Mouse Brain Atlas (see Methods) we quantified the anatomical distribution of tail-related vestibulo-spinal neurons among the subnuclei of the vestibular complex (Figure 3B). While neurons were found within all of the major (lateral, medial, superior and spinal) as well as accessory (x and y;^11^) subnuclei of the vestibular complex, they were predominantly residing within the spinal subnucleus (SPIV = 53.86 +- 7.75 %; Figure 3C) instead of the lateral vestibular nucleus as we had originally predicted (based on^10^). Intriguingly, we also found prominent labeling within nucleus X (X = 16.39 +- 2.88 %), one of the accessory vestibular nuclei that do not receive afferents from the vestibular organ (^11^). The spinal-projecting neurons in nucleus X were smaller than the other vestibulo-spinal neurons labeled in this study (VN_soma_ = 670.9 +- 17.54 µm, X_soma_ = 186.2 +- 10.29 µm; unpaired t-test, p*<*0.001; Figure 3D), indicating that they represent a morphologically distinct population within the vestibular nuclei.

**Figure 3:**
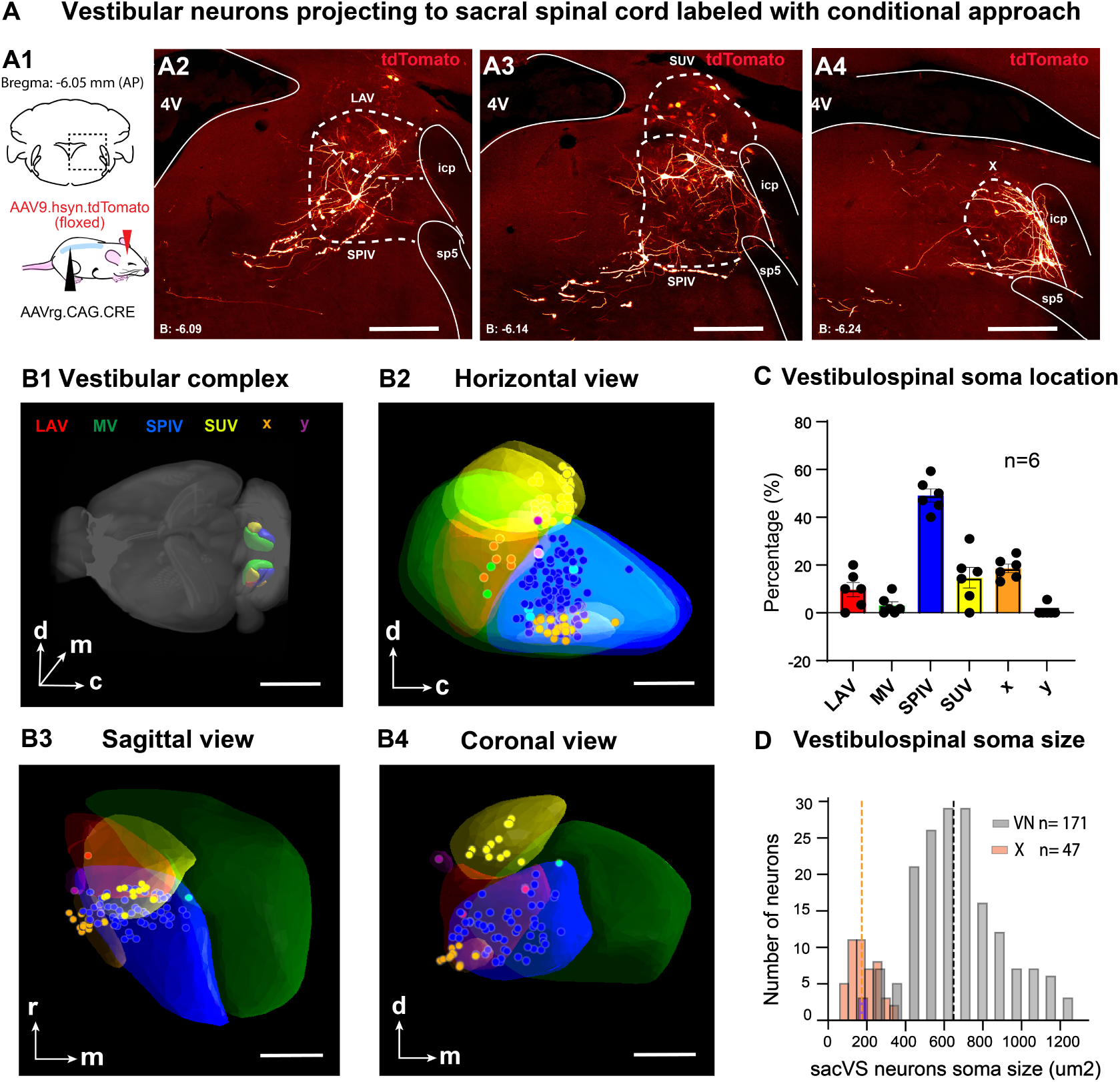
Vestibular neurons projecting to sacral segments are most commonly found in the spinovestibular nucleus. **A:** Retrograde intersectional viral labeling reveals vestibular neurons targeting sacral spinal segments. **A1:** Schematic of the viral strategy. **A2-A4:** Show example confocal images from a single animal with neurons labeled in different vestibular subnuclei. **B:** Localization of 218 neurons revealed by the conditional viral approach, registered to the Allen Mouse Brain Atlas (https://mouse.brain-map.org). **B1:** Overview of the vestibular complex in a 3D rendering of the mouse brain. **B2-4:** Close-up of vestibular subnuclei borders and the locations of identified neurons, shown from 3 different planes: Horizontal (B2), Sagittal (B3), and Coronal (B4). **C:** Distribution of neurons among the vestibular subnuclei. Each point represents data from a single vestibular nucleus (n = 6, from 3 mice). **D:** Comparison of vestibulo-spinal neurons’ soma sizes between main (grey) and accessory (orange) nuclei. Scale bars: A2-A4: 100 µm; B1: 3 mm; B2-4: 100 µm. Abbreviations: 4V, fourth ventricle; LAV, lateral vestibular nucleus; SPIV, spinal vestibular nucleus; SUV, superior vestibular nucleus; X, nucleus X; icp, inferior cerebellar peduncle; sp5, spinal trigeminal tract.

### Effect of vestibular nuclei activation on posture and tail movement

Successful balancing requires that the animal chooses the most context-appropriate motor strategy based on multimodal sensory perception (^18^). Thus, we wondered whether specific activation of vestibular complex neurons alone can trigger balance-related movements.

To examine this, we made use of targeted optogenetic stimulation of vestibular complex neurons by means of implanting an optic fiber coated with AAV9-hSyn-ChR2-mCherry-infused silk fibroin above the vestibular complex (Figure 4 A1; see Methods). This approach simplifies the surgical procedures and leads to ChR2 expression in neurons at close proximity to the optic fiber (Figure 2A2). Four weeks after the surgery, fiber-implanted mice were tasked with traversing either a flat surface (45 mm-wide) or the ridge (4 mm-wide) while their vestibular neurons were optogenetically activated using a 500 ms pulse at 1.5 mW (measured at the fiber tip, see Methods; Figure 4 A3).

**Figure 4:**
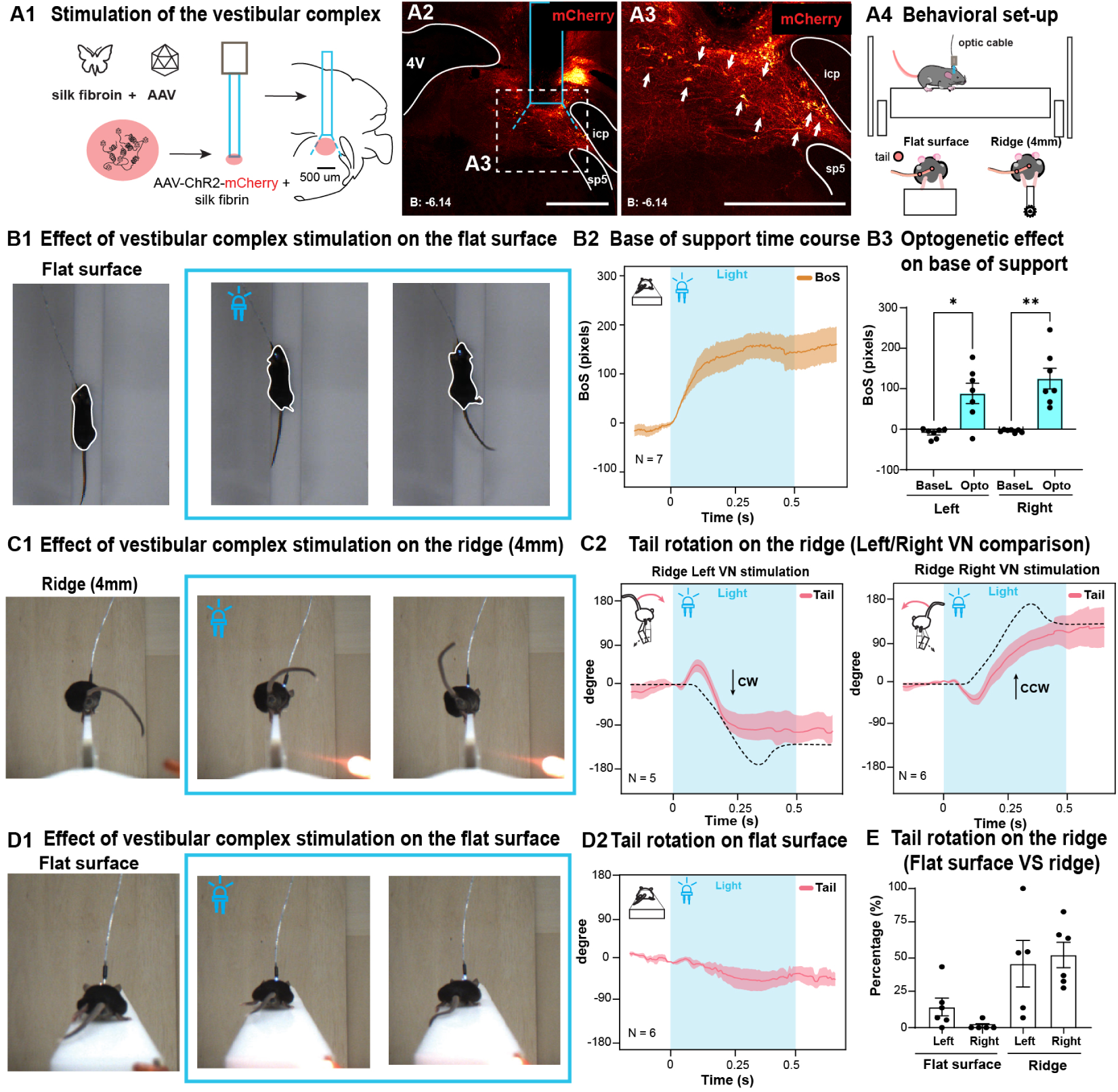
Postural balancing responses evoked by unilateral vestibular complex stimulation. **A:** Experimental design. **A1:** Strategy for localized ChR2 expression in the vestibular complex. **A2-A3:** Confocal image scan showing the location of the implanted fiber (blue rectangle) as well as mCherry fluorescence reporting on ChR2 expression in the vestibular complex. **A4:** Schematic description of the experimental set-up. **B:** Unilateral optogenetic activation of the vestibular complex leads to broadening of the base of support (BoS). **B1:** Single frames of a trial with optogenetic stimulation. The leftmost image is from locomotion preceding stimulation. **B2:** Changes in BoS quantified as pixel area for the mouse silhouette as seen from the top camera. Thick line represents the mean, and shaded area represents *±* SEM for VC photoactivation (N = 7). **B3:** Comparison of the BoS-change responses to left and right stimulation. **C:** Unilateral stimulation of the vestibular complex in mice on narrow ridges leads to countering body roll and tail swing responses. **C1:** Example frames from the rear camera taken prior (left) and during (middle and right) optogenetic stimulation. **C2:** Tail swing kinematics presented as change in angle (mean *±* SEM) in response to stimulation of the left (left plot, N = 5) and right (right plot, N = 6) vestibular complex optogenetic stimulation. Dashed lines represent the mean of tail-swing kinematics in response to a 20°tilt extracted from our previous work (^9^). **D:** Unilateral stimulation of the vestibular complex does not evoke body or tail roll responses in mice on broad ridges. **D1:** Example frames (as in C1) of VN optogenetic activation while the mouse crosses a flat surface. **D2:** Tail angle changes (presented as in C2) in the same mouse during flat surface crossing. **E:** Comparison of the percentage of stimulation trials during which a tail swing to the opposing side was observed, between broad and narrow ridges. Scale bars: A2-3: 500 µm. *, p *<* 0.05; **, p *<* 0.01; ***, p *<* 0.001. Abbreviations: 4V, fourth ventricle; LAV, lateral vestibular nucleus; SPIV, spinal vestibular nucleus; SUV, superior vestibular nucleus; X, nucleus X; icp, inferior cerebellar peduncle; sp5, spinal trigeminal tract.

As shown in Figure 4 B1, unilateral optogenetic stimulation of the vestibular complex led to a rapid postural adjustment including broadening of their base of support (BoS) by placing their paws further to the sides. We quantified the extent of the BoS as the area of the mouse silhouette seen from a camera above the ridge (Figure 4 B2) and showed that the BoS broadening is nearly identical regardless of the side on which the stimulation was delivered (Figure 4 B3).

In contrast, when the mice receiving vestibular stimulation were traversing the ridge (4mm), there were no changes in the BoS (supp Figure 4 A). The optogenetic activation elicited a slight tail rotation on the roll plane in the direction of the stimulation, followed by a larger rotation in the opposite direction (Figure 4C1). Intriguingly, kinematics of tail movement were similar to those described in our previous study evoked by roll-plane tilts of the ridge (see Figure 4 C2 for comparison of the mean tail swing responses with the responses to the tilts in^9^). However, no tail or body rolling responses were observed when mice were traversing the flat surface (Figure 4 D1-3). The optogenetic stimulation lead mice to a decrease in forward speed both on the ridge and on the flat surface, causing the mice to almost halt completely while swinging their tails (see supp Figure 4 B).

### Selective activation of tail-related vestibulospinal neurons elicits tail movement under certain conditions

In the previous section we described the effect of non-selective unilateral photoactivation of the vestibular complex. However the vestibular complex connects to several downstream targets which could elicit the postural motor responses we reported. To restrict optogenetic manipulation to vestibulospinal neurons projecting to the sacral spinal cord (see Figure 3 and 2) we injected retrograde-AAV vector (AAVrg.CAG.ChR-tdTomato) into sacral segments of the spinal cord (Figure 5 A). This allowed us to activate neurons targeting this region using an optic fiber implanted above the vestibular nucleus. As shown in Figure 5 B1-B2, optogenetic stimulation increased the likelihood of a tail swing during ridge crossing compared to non-stimulation trials. Importantly, the stimulation was effective in inducing tail swings only in trials where the stimulation was delivered to vestibulospinal neurons located on the same side as the tail position at the time of the stimulation (e.g. tail on the left, sacVS neurons on the left of the mouse activated; ”Ipsilateral trials”, Figure 5 B2). In ”contralateral trials” where the stimulation was delivered to the vestibulospinal neurons on the opposite side from tail position, no increase in tail movements was observed (Figure 5 C). This optogenetic stimulation of the sacVS pathway did not have any effect on the animals’ BoS (suppl Figure 5).

**Figure 5:**
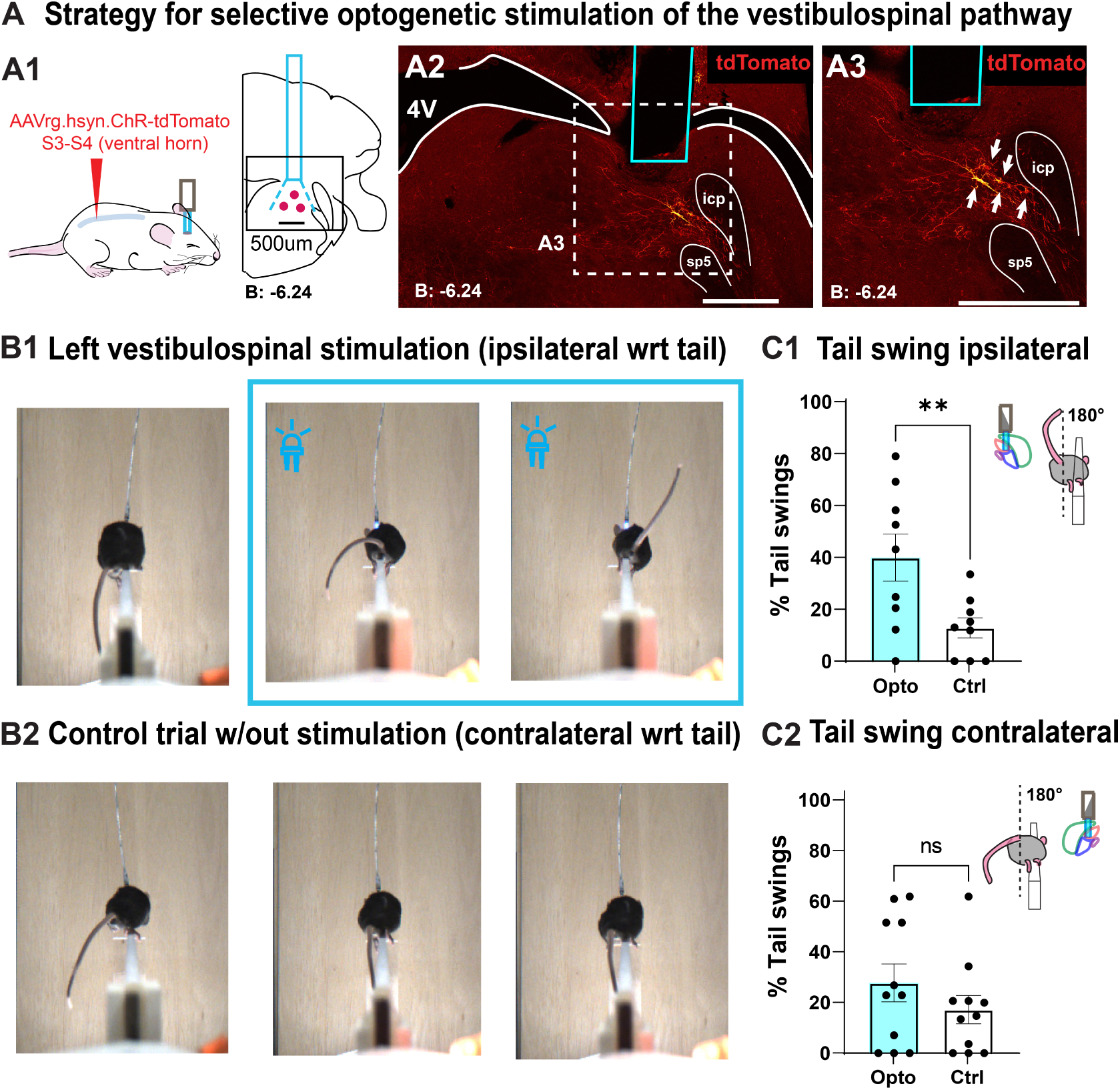
Tail movements evoked by unilateral stimulation of sacVS neurons. **A:** Strategy for obtaining selective expression of ChR2 in vestibular neurons projecting to the tail-related spinal segments. **A1:** Schematic of the approach; the blue rectangle represents the location of optic fiber placement in a separate surgery. **A2, A3:** Representative confocal images from the vestibular nuclei region showing retrogradely labeled neurons. A2: 10x magnification; A3: 20x magnification. **B:** Optogenetic activation of vestibular neurons targeting the sacral spinal cord increases tail swing probability only in ipsilateral condition. **B1:** Example frames from optogenetic stimulation during an ipsilateral trial; **B2:** Summary of all ipsilateral trials. Each dot represents data from stimulation on a given sacVS activation (N = 9, n = 17 ± 4 trials). **C:** Optogenetic activation of sacVS neurons on the opposite side of tail position fails to evoke tail movements. **C1:** Example frames from a contralateral trial; **C2:** Summary of all contralateral trials. Each dot represents data from stimulation on a given sacVS activation (N = 11, n = 15 ± 2 trials per animal). Scale bars: A2-3: 500 µm. *, p *<* 0.05; **, p *<* 0.001. Abbreviations: 4V, fourth ventricle; LAV, lateral vestibular nucleus; SPIV, spinal vestibular nucleus; SUV, superior vestibular nucleus; X, nucleus X; icp, inferior cerebellar peduncle; sp5, spinal trigeminal tract.

### Activation of sacVS neurons improve the selection of context-appropriate tail responses

Even though the ipsilateral sacVS stimulation increased the percentage of trials where mice swang their tails, the responses were not time-locked to the onset of the optogenetic stimulation (as they were for VN stimulation, Figure 4A supplementary), and occurred on average around 40 % of the ridge-crossings (Figure 5 C1).

As we have shown previously, mice swing their tails on the other side of the ridge when challenged with a tilt of the platform directed towards the side where the tail originally was placed (ipsilateral condition). On the other hand, tilting away from the tail (contralateral condition) results in a similar countering movement but no swing to the other side (Figure 6 A1). By using a strong perturbation (20 °tilt) we observed that ipsilateral trials reliably elicit a swing, and contralateral do not (^9^). By reducing the perturbation angle, we aim to approach a regime where the sensory stimulation by itself is not enough to reliably elicit a tail swing in the appropriate direction. Reducing the perturbation size from 10 degrees gradually decreases the percentage of tail swings in ipsilateral trials, reaching a point at the 5-degree condition where mice produce tail swings at similar rates, regardless of tilt direction (Figure 6 A2). We quantify tail-swing response accuracy as the fraction of correct responses (defining ”swings” and ”no-swings” as correct responses to ipsi- and contralateral tilts respectively) out of all trials (Figure 6 A3). The accuracy decreases from 0.66+-0.068 to 0.19+-0.043 (p=0.001, ANOVA) when decreasing the tilt magnitude, suggesting that the 5-degree tilt condition could be used as an ideal regime to examine if sacVS neurons activation leads to a performance improvement.

**Figure 6:**
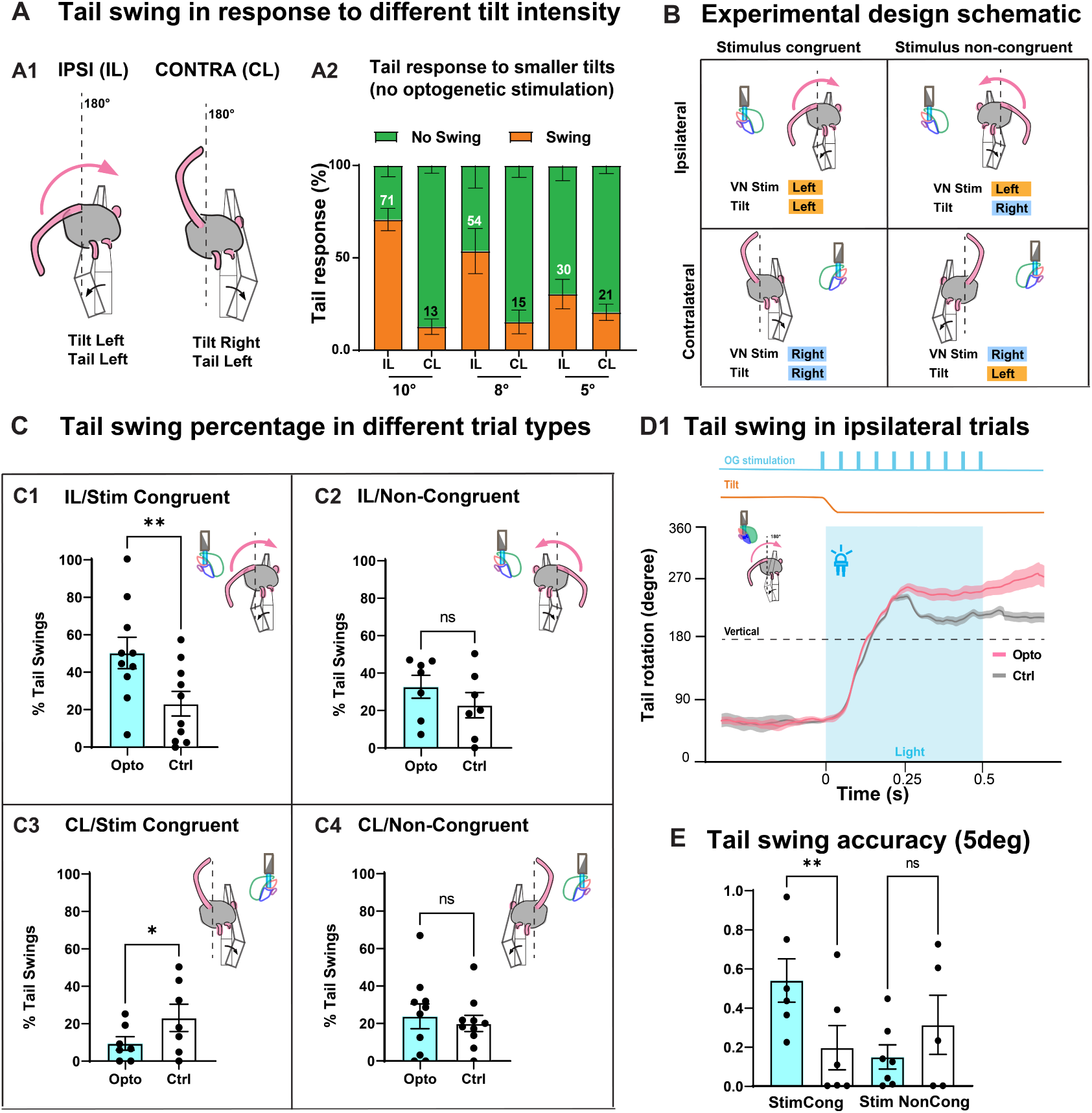
Context-appropriate activation of sacral-projecting vestibular neurons enhances the likelihood of tail engagement in response to roll-plane tilt perturbations. **A:** Definition of tail-swing response accuracy. **A1:** Definition of ipsi- and contralateral trials based on whether the tilt and tail are on the same side. **A2:** Percentage of trials where an appropriate tilt-countering tail-swing occurred decreases with smaller tilt magnitudes. **B:** Schematic description of the optogenetic experimental design. **C:** Percentage of tail swings observed in trial types as defined in B. **D1:** Tail swing kinematics in ipsilateral stimulus-congruent trials compared with those occurring in trials without optogenetic stimulation. Average angle trajectories for stimulation (red) and non-stimulation (black) trials; shaded areas indicate ± SEM. The blue shaded region indicates the time of optogenetic stimulation; the time of ridge tilt is indicated above the graph. **E:** Results from all trials compiled to show an increase in swing accuracy with stimulus-congruent trials (left) and no effect in stimulus non-congruent trials (right). *, p *<* 0.05; **, p *<* 0.001.

We constructed an experimental plan that examined tilt-triggered tail movements in four distinct contexts (Figure 6 B), defined by whether the stimulation was given on the same or opposite side as the tilt perturbation (”stimulus-congruent” or ”stimulus non-congruent”), and whether the perturbation was ipsi- or contralateral to the tail.

We expected to see that stimulus-congruent ipsilateral trials might have higher probability of a tail swing occurring in response to the 5-degree tilt, and indeed this was the case, rising from 12.81+-3.85 % of non-stimulated trials to 39.72+-8.97 % in stimulation trials (p = 0.0092, one-way ANOVA; Figure 6 C1). What we were more surprised to observe was that in the contralateral stimulus-congruent trials the percentage of tail swings decreased (from 22.97+-7.23 to 9.36+- 3.57; p = 0.023, one-way ANOVA;Figure 6 C2 ). Furthermore, the optogenetic stimulation did not lead to any increase or decrease in tail swings in non-congruent trials (Tail-Swing_IL/Opto_ VS Tail-Swing_IL/Ctrl_, p = 0.883; Tail-Swing_CL/Opto_ VS Tail-Swing_CL/Ctrl_, p = 0.556; One-way ANOVA. Figure 6 C3-4). The increase in percentage of tail swings in the ipsilateral stimulus-congruent trials was not associated with changes in the kinematics of tail movement(as shown in Figure 6 E).

In summary, vestibulo-sacral activation did not just drive tail movements in the direction opposing the stimulus, but rather improved the accuracy of action selection (swing or no swing) when delivered congruently with the tilt perturbation, without affecting its kinematics.

## DISCUSSION

In this study we provide the first detailed description of the neuro-muscular organization of the mouse tail, and show that the tail motoneurons in the spinal cord are directly targeted by a subpopulation of vestibulospinal neurons in the brainstem. These findings expand previous research on the importance of the mouse tail in fundamental aspects of balance and posture by introducing the neuronal substrates responsible for the tail control in balance.

### Anatomical insights into tail neuro-muscular control

Our high-resolution microCT-based reconstruction of the proximal and mid regions of mouse tail evokes evolutionary considerations.

The tetrapod vertebral column is a classic example of segmented body organization at a structural level, showing regional diversification to support and allow optimal motility (^19,20^). Besides skeletal specializations that define the preferred directions of movements, organization of muscles and tendons with respect to the segment boundaries expands motor capabilities by providing support for speed, strength or motility range for increased ecological fitness (^21^).

Metamericity and division into extrinsic and intrinsic tail musculature has been previously proposed^16^ based on observations during manual dissection. Specifically, the basal (most proximal to hips) extrinsic muscles span multiple vertebrae, while intrinsic muscles operate over single vertebral joints distally. This organization suggests a division of labor in tail control, so that the extrinsic muscles are positioned to generate large movements, while the intrinsic muscles control finer motions such as tail coiling. Here we extend the earlier observations by revealing presence of long tendons inside (as well as between) the extrinsic lateral muscles and hence we classify them as bipennate rather than bicepetal muscles (^22^). This structural organization may play a key role in storing elastic energy, facilitating rapid, propulsive tail movements originating from the base (shown previously in^9^). Parallels to such arrangement can be found for example in the hindlimb tendons of animals like kangaroo rats, known for their fast and propulsive jumps during their escape response (^17^).

### Localization of tail-controlling motoneurons in the sacral-coccygeal spinal segments

We characterize the distribution of tail-motoneurons (tail-MNs) in the sacral-coccygeal spinal segments (S3-Co3) using retrograde labeling. We found that tail-MNs are distributed caudally to the hindlimb-motoneurons, which reside predominantly in lumbar segments (T13-S1;^23^). We observed a topographic organization, with the distal (intrinsic)-targeting motoneurons found more caudally with respect to those targeting the proximal muscles (S3-4 vs. S4-Co3). Furthermore, the extrinsic - and intrinsic-targeting motoneurons differed in soma size, suggesting a division of labor based on Henneman’s principles^24^ where extrinsic motoneurons (larger in size) would require a stronger input in order to recruit extrinsic muscles, compared to the recruitment of intrinsic motor units.

Despite their distinct rostro-caudal distribution, both proximal- and distal-targeting motoneurons formed a localized ventro-medial clusters within the spinal laminae VIII and IX, putatively corresponding to the medial motor column, similarly to what reported in axial muscles (^25^). Similar to the tail, hindlimb motoneurons are also organized along the rostro-caudal axis. However, the hindlimbs motoneurons targeting proximal (extrinsic) muscles are found more medially, while those innervating distal (intrinsic) muscles are located more laterally (as well as more caudally) (^26^). This organization is preserved across different species, such as mice (^26^), birds (^27^), reptiles (^28^), and non-human primates (^29^). This may be important for forelimbs control due to the evolution of dexterity, where control of intrinsic muscles may require selective supraspinal input (^30,31^). In contrast, the organization of tail motoneurons in one localized medial cluster may reflect the fact that different tail muscles recruitment rely mostly on local spinal cord circuitry, rather than different descending supraspinal pathways.

### Descending vestibular pathways to tail-related spinal segments

Previous literature has described two main descending vestibular pathways that influence spinal motor programs. One pathway originates from the medial vestibulospinal nucleus and sends axons primarily to the cervical spinal cord, where it helps stabilize the head via the vestibulo-collic-reflex (VCR^32^). The second pathway originates from the lateral vestibular nucleus and extends throughout the spinal cord where it contribute to postural adjustments (e.g. by activating hindlimb motoneurons^10^).

In this study, we identified a third descending vestibular pathway that originates predominantly from the spinal vestibular nucleus (SpVN) and targets mostly the intermediate layers of the spinal cord (lamina VII-VIII; Figure 3B1-B2). A previous study showed that injection of a rabies construct in the lumbar spinal cord led to sparse labeling in SpVN (^33^). However the work from Murray’s lab did not reveal any spinal vestibular neurons after fluorogold injection in the lumbar spinal cord, nor via transynaptic labeling from hindlimb muscles (which instead led to labeling of neurons in LVN^10^).

One intriguing explanation is that while the vestibular pathway originating from LVN forms direct projection to spinal motoneurons, the SpVN may project more dorsally to intermediate layer of the spinal cord (where interneurons are located). Indeed we found using our AAV-based retrogrde labeling that the axons originating predominantly from SpVN project mostly to lamina VII and lateral portion of lamina VIII (3 B1-B2), while our rabies-based transynaptic strategy labeled exclusively neurons in the lateral vestibular nucleus, indicating that the LVN originating pathway project directly to tail motoneurons (located in the medial portion of lamina VIII and IX, 1 F). The different output of these two pathways raise questions on whether different subnuclei of the vestibular complex process different information, as they target different output system in the spinal cord. Given that the vestibulospinal tract runs mostly ipsilaterally in spinal cord segments caudal to the cervical spinal cord (^34,35^), the LVN-originating part of the vestibulospinal tract can only control tail muscles on the same side. However in order to perform a full tail swing the sequential recruitment of muscles from both ipsilateral and contralateral sides are necessary. Hence we expect that the SpVN originating portion of the tract plays a major role in tail swing, given that the recruitment of the interneuron layer in the spinal cord is more likely to have a more widespread downstream effect in recruiting different muscle groups both ipsilaterally and contralaterally.

### Vestibular nuclei provide sensory context for appropriate tail-response

One striking aspect of our functional analysis is that photoactivation of the vestibular nuclei (VN) resulted in tail rotation only on the more challenging balancing condition. This rotation mimics the tail motor response we have previously reported in response to an external, roll-plane perturbation (^9^). We noted an important difference in the two responses. In particular, during optogenetic stimulation the first tail rotates slightly towards the ipsiverse side before swinging to the opposite side, unlike in the tilt-induced trials (see full vs. dashed lines in 4 C2). One possible explanation for the observed phenomenon is that the early phase of the response (ipsilateral rotation) is due to activation of the direct pathway from vestibular nuclei to motoneurons, while the later phase (contralateral rotation) can be attributed to the recruitment of the interneuron circuit in the sacral spinal cord (in particular commissural interneurons which allow the activation of motoneurons in the contralateral side to the photo-activated circuit).

On a flat surface no significant tail movements were elicited with the optogenetic approach, but we observed a widening of the base of support, similarly to what has been previously described (^36^). One mechanism that can explain the tail rotation being evoked on the ridge but not on the flat surface relates to the the differences in hindlimb proprioceptive signaling during flat surface versus ridge-crossing. A narrower stance requires greater muscle tension to maintain the mouse body upright (^37,38^), which possibly enhances proprioceptive sensory feedback to the motoneurons (^39^). A possible substrate for the relay of hindlimb proprioceptive information to the tail spinal circuits is propriospinal interneurons in the spinal cord (^40^). Indeed, our transynaptic tracing from the tail labeled interneurons in the lumbar spinal cord (3 supplementary), which indicates that these lumbar interneurons contact directly tail motoneurons. While more work is needed to examine the identity of these interneurons and their potential role in tail control, it is plausible that they could work together with the vestibulospinal input in providing sensory information (particularly from the hindlimbs) for tail control.

Here we investigated whether the direct sacVS pathway can evoke tail swings by restricting the optogenetic stimulation to vestibulospinal neurons projecting to the sacral spinal cord (5 A). At first we were puzzled by the results obtained with the restricted stimulation. No BoS-related postural adjustments were observed (Supplementary Figure 4), but the tail responses elicited were not time-locked to the stimulation, suggesting that sacVS stimulation by itself is not enough to reliably induce tail rotation across conditions (although the responses evoked retained the dependency on context, as we observed an increase in tail rotation only when the tail was positioned ipsilaterally to the stimulation). One possibility is that the sacVS pathway increases the likelihood of tail rotation (under certain sensory contexts). Another is that the sacVS pathway informs the spinal cord of the direction of the response, increasing or decreasing the likelihood of a tail rotation depending on the sensory context. This set us out to investigate whether priming the sensory system (by inducing a small tilt at the onset of the optogenetic stimulation) would lead to discriminate between these two possibilities.

We found that by using a 5°tilt we evoked the same percentage of tail swings regardless of the direction of the tilt. This showed that under the 5 °tilt condition mice do not produce a directional-appropriate response (as they did with larger tilt amplitudes, 6 A3). The optogenetic activation of the sacVS pathway leads to an increase in the accuracy of eliciting a tail swing under the appropriate sensory context (i.e. increase in tail swing in ipsilateral, and decrease in contralateral). This modulation of the tail swing response suggests that the sacVS tract informs the spinal cord about the direction of the perturbation, allowing the spinal circuitry to generate the appropriate motor response up to what the same mice showed for larger tilts (10 °, 6 A2). While we observed this increase in accuracy in conditions where the stimulation side matches the tilt direction (stimulus-congruent trials), we did not report the same effect when the direction of the activation is opposite to the perturbation (stimulus non-congruent condition, 6 C2-C4). Taken together, these results indicate that the sacral-vestibulospinal pathway provides information about the perturbation direction to the spinal cord, enabling more accurate postural adjustments to the sensory perturbations, thus enhancing the mouse’s ability to maintain balance under challenging conditions.

### Limitations of the study and future directions

An unexpected finding in this study was the significant Nucleus X (NuX)-originating axonal projections to the tail-related spinal cord segments. To our knowledge, NuX was not known to project to the spinal cord in mice or other species. As NuX does not receive afferents from the vestibular organ^41^, it remains to be understood how it contributes to tail movement. One possibility is that such contribution may originate from the spino-cerebellar pathway, which relays proprioceptive information originating from the lumbar spinal cord to nucleus X, as shown by single axonal morphology reconstruction (^42^). Unfortunately, as to our knowledge there is no genetic tools to selectively target distinct vestibular sub-nuclei, we were not able to dissect the different functional contribution of nuX and SpVN, and thus we can not speculate on their specific roles. We expect that further genetic profiling of vestibular neurons (such as in^43^) will lead to elucidation of the functional roles of distinct vestibular sub-nuclei.

In our study we focused on the role of the vestibular system in tail control, but it is likely that other neuronal circuits, providing proprioceptive and exteroceptive information, can play a role in the tail-swing response. For example, the function of hindlimb spinal circuits in modulating tail movement is not known, but our previous results^9^ as well as prior studies^44^ shows that the tail oscillations are phase-locked to the hindlimb step-cycle during locomotion. Functional manipulation of lumbar spinal interneurons projecting to the sacral spinal cord can reveal important insights into the relation between the hindlimbs and the tail.

### Tale of mammalian tails

Throughout animal evolution, balancing and maintaining context-appropriate posture have remained essential components of movement and behavior. The transition to terrestrial life and legged locomotion introduced a new challenge: maintaining the center of mass over the base of support during movement. Given that the vestibular system, one of the most ancient sensorimotor structures, plays a key role in postural control (^45^^–47^), its function had to adapt to the increasingly complex demands of balancing from swimming to terrestrial locomotion (^48^). This evolutionary shift parallels the emergence of more complex body plans, which likely spurred the refinement of balancing strategies. In chordates, this is particularly reflected in the evolution of tails as flexible and muscular structure capable of generating compensatory angular momentum to stabilize the body during locomotion on narrow substrates (^5^). Given the evolutionary significance of tails in maintaining balance across species, studying tail dynamics in mice provides an ideal model for investigating the neural mechanisms underlying postural control.

Studying tail movements in mice offers unique advantages for understanding neuromechanical control strategies. Unlike limbs, the tail provides a simpler, more tractable system for investigating balance, allowing us to isolate the contribution of the vestibular system to postural control. The tail is an ancient evolutionary innovation omnipresent across different clades of the tree of life. Understanding the neuronal control of tails in mice, leveraging the tools for genetic and neural circuits manipulation available in this species, may shed light to fundamental principles of design of neuronal circuits for locomotion that are shared across species.

## Acknowledgments

We would like to acknowledge Dr. Shinya from the OIST Imaging Section, and the OIST Animal Facility for their technical support. We would like to thank Dr. Fumiyasu Imai for the help in the design of the rabies experiment. We also would like to thank professor Graziana Gatto for helpful feedback on the manuscript. S.A.L. is supported by JSPS DC1 fellowship (202020494). N.I. and M.Y.U. are supported by OIST intramural funding.

## Author contributions

Conceptualization, S.A.L. and M.Y.U.; methodology, S.A.L., N.I, and H.H.; investigation, S.A.L. and M.Y.U.; N.R. transynaptic tracing; J.K. tail muscle reconstruction; writing – original draft, S.A.L. and M.Y.U.; writing – review & editing, S.A.L., M.Y.U.; funding acquisition, S.A.L. and M.Y.U.; resources, M.Y.U., and Y.Y.; supervision, M.Y.U.

## METHODS

### EXPERIMENTAL MODEL AND SUBJECT DETAILS

#### Mice

All procedures were reviewed and performed in accordance with the OIST (Okinawa Institute of Science and Technology) Graduate University’s Animal Care and Use Committee (protocol 2020-292-2). The mice (C57BL/6, male, 6-12-weeks-old) were purchased from Clea (Tokyo, Japan) and were acclimatised to the OIST animal facility for at least 3 days before handling or surgery. They were housed in institutional standard cages (maximum of 5 animals per cage) on a reversed 12-hr light/12-hr dark cycle with ad libitum access to water and food.

### METHODS DETAILS

#### Micro-computed tomography (µCT) and 3D visualization

The tail was carefully dissected from an adult mouse (C57BL/6, male, 12-weeks-old), after transcardic perfusion of the animal with 4% paraformaldehyde (PFA, Electron Microscope Sciences, Pennsylvania, United States). Upon dissection, the tail was fixed in 4% PFA at 4°C for 24 hours to preserve the tissue. Subsequently, the tail was dehydrated through a graded series of ethanol solutions (30%, 70%, and 100%) and stained using an iodine potassium iodide (I2KI) solution (Lugol’s solution: 1% I2, 2% KI in distilled water) for 48 hours to enhance soft tissue contrast for micro-CT scanning. Following staining, the tail was rinsed in 100% ethanol and air-dried to wash out the excessive iodine. The I2KI-stained tail was scanned with a Zeiss Xradia 510Versa µCT system, operating at 50 kV, 79 µA, and 4 s exposure time with a 0.4 X magnification lens. The resulting data was reconstructed with an isometric voxel size of 16 µm3 using Amira software version 2020.2 (ThermoFischer Scientific, Massachusetts, USA). We cropped the µCT dataset at the edges of the field of view (FOV) and filtered the image using Amira’s 3D Bilateral Filter (similarity: 5000) for further processing. We segmented the tail muscles using Amira. Briefly, we first created pre-segmentations in all three planes that we then interpolated using the Biomedisa toolbox (^50^). We exported individual muscles as well as vertebral bones as STL files. We then proceeded to import and visualize those using Blender 4.2 (Blender Foundation, Amsterdam, Netherlands).

To highlight tissue structures (such as the lateral tendon in yellow in Figure 1 A) beyond the muscles, we used another µCT scan of the same specimen at higher resolution (40kV, 74 µA, 10 s, 4 X, 4.8 µm3) to create a volume rendering of selected muscles and tendon. In Amira, We masked and partitioned the µCT scan into 4 parts and with differently colored transfer functions in Drishti 3.2 (^51^), we highlighted a dorsal extrinsic muscle (blue), a lateral extrinsic muscle (red), the 4th coccygeal vertebra (white) and the other vertebrae (dark grey).

#### Surgical procedures for anatomical tracing

##### Tail injections

The animal was first anesthetized with 3-4 % isoflurane in an induction chamber using the nosecone connected to an anesthetic system (SomnoSuite, Kent Scientific, Connecticut, USA). The hair was removed using hair-remover cream, and the area was disinfected with iodine and anesthetized locally using lidocaine (Sandoz, Basel, Switzerland). Then the skin was removed carefully using a curved spring scissor. The tail was made taut using a spinal cord clamp (customized device N0-008-004 from Narashighe, https://www.narishige.co.jp, Japan), and a small portion of the muscular fascia was removed to facilitate the insertion of quartz glass pipette (Sutter Instrument, California, USA). A total volume of 1 µl or 700 nl was injected on each extrinsic or intrinsic muscle respectively at a rate of 150 nl/minute. The pipette was then gently retracted, with a pause of 30 seconds before the pipette tip was removed from the muscle. Post-surgical care included closing the injection site with suture, applying topical anesthetic, and administering systemic analgesics such as carprofen or buprenorphine.

##### Spinal cord injections

For spinal cord injections, after induction of anaesthesia and application of topical analgesic, the fur along the back of the mouse was shaved, and the skin cleaned. The skin and fascia covering the lumbar spine (L1-L3) were carefully opened. A small slit of the caudal portion of the L2 coccygeal bone was removed dorsally to expose the caudal portion of the sacral spinal cord, and the same viral constructs were delivered as described above. 500 nl of virus was injected at a rate of 50 nl/min. After the injection, the pipette was withdrawn slowly, and both fascia and skin were sutured. Post-operative care included administering systemic analgesics and closely monitoring the animal for several days to ensure full recovery.

##### Stereotaxic surgery in the vestibular complex

For brainstem stereotaxic brain injections, the hair on the scalp was removed and the skin was cleaned. An incision was made in the skin to identify bregma and lambda. A small opening (*<*1 mm diameter) was made using a surgical drill, and the viral constructs were injected stereotactically at coordinates AP -6.20 mm, ML +-1.55 mm (from Bregma), and 30 nl of the virus was injection at each depth of DV -4.5 and -4.4 mm. As with the tail and spinal cord, a quartz-glass pipette was used to deliver the virus, but for brainstem injections we delivered at a rate of 10 nl/min into the vestibular complex. After the procedure, the skin above the skull was closed with sutures. For postoperative pain management, 0.05 mg/ml Rimadyl (Zoetis, New Jersey, US) was administered subcutaneously at a dose of 5 mg/kg, and the mouse was monitored regularly during recovery.

#### Viral constructs

The virus constructs used were adeno-associated virus (AAV) purchased from Addgene (Massachussets, USA), with capsid AAV9, or AAV retrograde, and promoters were either hsyn or CAG (as specified in our virus transfection protocol RDE-2022-041-2, with titer *≥* 7 *×* 10^12^ vg/mL). Four to six weeks of labeling/transfection time were needed after the injection of viral constructs. During this post-operative phase, the animals were regularly monitored to detect any possible problems in recovery.

#### Rabies tail muscle injection

For transsynaptic labelling via intramuscular tail injection, N2cdelG-tdTomato rabies virus (titer 1X108, Center for Neuroanatomy with Neurotropic Viruses (CNNV), Pennsylvania, USA) was complemented with equal volume of AAV6-oG-P2A-H2BStayGold. 5-day old C57BL/6J pups were anesthetized by placing them on a glass cover over crushed ice (so to prevent the pup’s skin to be in direct contact with ice) for around 3-5 minutes. 4 µl of mixed virus solution (2 µl rabies+ 2 µl AAV6) were injected at each site, 1mm from the tail base: dorsal muscle injection D/V: - 0.3mm and lateral muscle injection D/V: -0.3mm and -0.6mm. 8 days post virus injection, pups were perfused transcardially using ice-cold PBS and 4 % PFA. Brain, spinal cord, and tail tissues were harvested. Upon dissection, brain, spinal cord and tail tissues were fixed additionally at 4 C°in 4% PFA before transfer to 30% sucrose. 50 µm cryosections of brain samples were counterstained with NeuroTrace 500/525 to label neurons and imaged using Leica SP8 confocal microscope. ChAT Ab was used to label motor neuron pools in sacral spinal cord sections.

#### Optic cannula preparation coated with silk fibroin

The optic cannulas were prepared using a 100 cm (length) by 400um (diameter) optic fiber (Thorlabs, New Jersey, United States), that was cut using a ruby cutter (Thorlabs, New Jersey, United States) to obtain 1 cm long optic fiber. The fibers were glued to a metal cannula (Thorlabs, New Jersey, United States) and then they were disinfected with ethanol (70 percent). The coating method was adapted from (^52^). The cleaned implants were positioned vertically and the Nanojector (Neurostar, Tubingen, Germany) positioned above the implant. 150 nl mixture (1:1 mixtures of silk and virus mixed with a pipette) was applied by slowly dispensing in 25 nl droplets on the face to be implanted, letting it dry before another droplet was added. Fibers were then vacuum-desiccated overnight at 4 °C and then used for implantation within one day.

#### Surgical procedures for implants in the brainstem

At the beginning of the fiber implantation surgery, anesthesia was induced with 3-4 % isoflurane and then maintained at 1.8 % isoflurane throughout the procedure. The head was secured using mouse ear bars, and pre-operative measures included the application of eye ointment, scalp shaving, and lidocaine gel application for topical anaesthesia. An incision was made from between the ears to between the eyes, and the skull was cleaned of periosteum before being dried. A black enameled 0.4 mm needle was installed in the stereotaxic apparatus (Kopf, California, USA). The brain was leveled by aligning Bregma and Lambda, and the alignment on the dorso-ventral axis was checked not to exceed 50 µ. The needle was moved to the midpoint avoiding values larger than 50 µm. Craniotomy was performed with a drill (NSK, Tochigi, Japan) to reach the coordinates (AP 5.9, ML +-1.55, DV 4.35) with the implant slowly placed using the stereotactic robot (Neurostar, Tubingen, Germany). After implant placement, a small drop of super glue was applied at the insertion point on top of the skull to fix the implant in place, left to cure for 5 minutes, and the implant was released from the stereotaxic robot. The second implant was installed on the opposite side following the same procedure. Once cured, dental cement (Super-Bond, Sun Medical, Moriyama, Japan) was mixed according to product use manual in a ceramic dish and applied to the skull and implants. The skin was pulled over the cement, stitches were placed in front and behind the implant, and the mouse was returned to the cage on a heat pad. For postoperative pain management, 0.05 mg/ml Rimadyl (Zoetis, New Jersey, US) was administered subcutaneously at a dose of 5 mg/kg, and the mouse was monitored regularly during recovery.

#### Histology

##### Perfusion and slicing

The brain and spinal cord of the mice used for anatomical investigation was collected after perfusion-fixing. The animal was deeply anesthetized with MMB (containing Medetomidine HCL, 1 mg/ml; Midazolam, 5 mg/ml; and Butorphanol tartrate, 5 mg/ml; dosage 0.05 ml/g) and after they were deeply anaesthesized (verified with an absence of the toe pinch reflex), the heart was exposed and the animal was transcardially perfused, first with phosphate-buffered saline (PBS; pH 7.4) to clear the blood and then with fixative solution (4 % paraformaldehyde, PFA), and finally the tissue was extracted and left in 4 % PFA overnight in a fridge at 4°C . After fixation, the tissue was immersed in 10 % sucrose in PBS until it sank (6-12 hours), followed by immersion in 20 % sucrose for another 6 hours, and in 30 %sucrose in PBS overnight for cryoprotection. Coronal brainstem sections, 50 to 100 µm thick, were cut using a vibratome (5100MZ-plus; Campden Instruments, Loughborough, UK) with ceramic blades (38 x 7 x 0.5 mm, model 7550-1-C, Campden Instruments, Loughborough, UK) or a Cryostat (Leica CM1950, Wetzlar, Germany). The sections were then mounted on slides with Vectashield (H-1200, VectorLabs, MA, USA) mounting medium and covered with 1.5 coverslip glass (Harvard Apparatus, MA, USA)

##### Image acquisition

Confocal image stacks were obtained from sections labeled using viral methods with a Zeiss LSM 880 confocal system (Zeiss, Germany). Low-magnification imaging utilized a 5x objective (Plan-Apochromat 5x M27; NA 0.16; Zeiss, Germany) with z-steps between 3 to 8µ. High-magnification imaging was performed with a 40x objective (Plan-Apochromat 40x Oil DIC M27; NA 1.4; using Zeiss Immersion oil; Zeiss, Germany) with z-steps between 0.1 to 1 µm for tissue sections between 50 to 100 µm thick. For multi-channel images acquired in line-scan mode, the following excitation/emission wavelengths were used: eGFP (and AF 488) at 488 nm / 490–535 nm, and tdTomato at 561 nm / 470–655 nm.

##### Pre-processing overview

Image pre-processing involved applying a Gaussian blur (1 pixel), before optimizing brightness levels to enhance the visibility of neuronal soma outlines. Then we traced neurons in ImageJ. Traced neuron silhouette somas were saved in the ROI manager and z-projected. In case of overlapping neurons we divided the stacks so that each neuron will be located in a given substack. For further processing and analyses, custom scripts were used (Python 3.7.4; https://www.python.org run on Windows 10).

#### Spinal cord registration onto the Allen atlas and density map reconstruction

Prior to the registration step, we identified the identity of the spinal cord region based on the length of the horizontal line that divides the dorsal horn from the rest of the spinal cord. The measurements were obtained using the Allen Spinal Cord Atlas https://mouse.brain-map. org/experiment/siv?id=100050403&imageId=101006559&imageType=ref_series. The table 1 provides the criteria that we used to determine the spinal cord segment based on line length measurements (unit is micrometer).

**Table 1:** Line length thresholds for different spinal segments.

| Spinal Segment | Line Length Threshold ($\mu\text{m}$) |
| --- | --- |
| S1 | < 1600 |
| S2 | < 1300 |
| S3 | < 1150 |
| S4 | < 950 |
| Co1 | < 800 |
| Co2 | < 700 |
| Co3 | < 650 |

After deciding the spinal segment identity, we proceeded with the registration of the confocal acquired data into the atlas, using the ’Align Image by Line ROI’ toolbox in Fiji to allow rigid transformation, and alignment of the image onto the atlas. After quality check, if the image registration resulted distorted on the dorso-ventral axis, the procedure was repeated by drawing a second ROI that passes vertically by the central canal (CC). These steps were repeated manually for each stacks obtained, and a mask containing the area of interest (either segmented axons or soma location) was extracted and data for the same mouse and same segment were overlaid with z-projection. Spinal cord registration onto the Allen atlas and density map reconstruction.

#### Spinal cord axon density map reconstruction

The spinal cord section was de-noised in Fiji using the background subtraction macro (http://microscopynotes.com/imagej/index.html). Then a threshold was applied to identify labeled axons in a binarized image, and any remaining noise, identified as scattered pixels, were deleted. We applied the rigid transformation described above to register the image on the spinal cord atlas. Finally, we used a custom-made script written in Python (Python 3.7.4; https://www. python.org run on Windows 10) to generate a density map and plot it on top of the atlas (obtained from^53^).

#### Brainstem registration on the Allen Atlas and 3D projection of vestibulospinal neurons in the Common Coordinate Framework (CCF)

We performed the registration of brainstem sections data onto the Allen Atlas (mouse.brain-map. org) using ABBA (Aligning Big Brain Atlases, EPFL, Switzerland,^54^). After registration we proceeded to manual localization of labeled sacVS neurons using QuPath (University of Edinburgh, United Kingdom,^55^), and projection of these coordinates in the Common Coordinate Frame-work (CCF), visualized in 3D using BrainGlobe (Sainsbury Wellcome Centre, UCL (UK), and the Max Planck Institute for Neurobiology (Germany),^56^). Specifically, we loaded the sections in ABBA, and after aligning each section to the respective Allen Atlas coronal section along the rostro-caudal axis, we performed linear registration, followed by non-linear registration (similar to what is described in^54^). Then the registered sections were loaded in Qupath and the labeled vestibulospinal neurons were detected manually. The 3D points (corresponding to the location of the somata) were transformed into CCF coordinates using a custom-written script in Python. The coordinates were then visualized in either 2D or 3D using the Napari toolbox in BrainGlobe (https://github.com/brainglobe/brainreg-napari).

#### Behavioral experiments

##### Animal training

After a 2-week handling and habituation period, mice were trained to cross the platform using the 4 mm ridge. The mice were gently encouraged to traverse the ridge, and only providing additional incentive for traversing by placing a familiar tube mice use as shelter in their home cages in front of them if necessary. No food or other rewards were used, and the animals were not incentivised artificially (e.g. with food deprivation or stressors). Each training session lasted until the mouse could cross the platform without stops for 10 times in a row. Each training session lasted for approximately 2 hours. Each mouse underwent such training for five consecutive days. All mice used in this study successfully acquired the task and traversed the ridge at the end of the 5th training day. No prior training was necessary for the flat surface crossing.

##### Optogenetic stimulation in the ridge task

Behavioral experiments started 21 days after the viral injection and/or optic fiber implantation. Implanted optic fibers were connected to an LED source (473 nm DPSS system, LaserGlow Technologies, Toronto, Canada) through a mating sleeve (Thorlabs, New Jersey, USA). For stimulation of the entire vestibular nucleus we used a continuous pulse (duration 500 ms) at 1.5 mW. In case of selective sacVS activation light was delivered in trains of pulses of 15 ms pulse width at 20 Hz frequency for a duration of 500 ms (values based on the vestibulospinal stimulation used in Murray et al.^10^). Prior to the experimental recording, in a pilot study we examined the responses and found the minimal laser power sufficient to evoke a response, which was measured to be between 5-10 mW (measured at the fiber tip using a power meter (PM100USB with S120C silicon power head, Thorlabs, New Jersey, USA)), to restrict photo-activation (^57^), prevent heat, and exclude an unintentional silencing by over-activation (^58^).

##### The behavioral tasks

A diagram illustrating the ridge task is shown in Fig. 4 A4. The ridge set-up consists of a thin acrylic ridge (4mm wide, 50cm long) attached on one end to a motor (S3003 Servo Motor, Futaba, Japan) controlled by a Bonsai script (https://bonsai-rx.org/) that can tilt the ridge left and right direction with varying amplitudes. A small platform was placed at both ends of the ridge for the mouse to comfortably reside before and after the trial. Mouse movement was recorded at 300 frames per second (FPS) with two high-speed cameras (Blackfly S USB3, Tele-dyne FLIR, Wilsonville, OR, USA), one placed 50 cm above the ridge and another at the rear end of the ridge. Video recordings were saved in mpg format. The plat surface in place of the ridge when necessary to record, using the same cameras used in the ridge. This surface consists of plexiglass as well and it is 45mm wide, 50 cm long and 9 cm tall.

##### Tilting ridge trial structure

Following animal training (see above), during the experimental recording mice were subjected to trials with a tilt perturbation randomly intermingled with non-perturbation trials, and the onset of the tilt was randomly timed during a trial to prevent anticipatory behavior. After the tilt, the ridge remained in that position for 3 second before returning to the original orientation. The perturbation trials involved a 10, 8 or 5 degree tilt of the ridge in either left or right direction. In addition to the 4mm ridge we used a 45-mm wide surface that allowed comparison of ridge-traversing locomotion to ”flat surface locomotion”. In additional trials the perturbation angle was either decreased or increased (to 10 or 30 degrees (150 or 230 ms), respectively) without changing the angular velocity of the tilt.

### QUANTIFICATION AND STATISTICAL ANALYSIS

#### Pose estimation and quantification of body using DeepLabCut and Bonsai

Videos were recorded with the top and rear cameras as described above. The tail trajectories were extracted using DeepLabCut (version 2.1.7; https://github.com/DeepLabCut/DeepLabCut^59^ A total of 200 image frames (10 videos selected from perturbation/non-perturbation trials, as well as different widths, 20 frames/video) were used to label and train models from both camera views. 90% of the labeled frames were used for training, and the remaining 10% for testing. We used a ResNet-50-based neural network for 1,000,000 training iterations, where the cross-entropy loss plateaued to 0.001. We then used a p-cutoff of 0.9 to condition the X,Y coordinates for future analysis. This network was then used to analyze videos.

In addition to the tail rotation kinematics, position of the body centroid was estimated using the top camera view as the average position of the mouse body silhouette using a custom-made script in Bonsai. Silhouette was extracted by first applying a filter to binarize the image (to separate the region of interest (ROI) from the background), and then extract the ROI centroid, as described in (^60^). The silhouette area was used as a proxy for base of support (BoS) measurement used in 4. The point in time were the ROI centroid became visible under the top camera were used to extract the time-aligned traces captured by both cameras. Out of this trace the 500 points (centered in time) of the time series were used to compute angles changes during a trial.

#### Kinematics analysis

Custom Python scripts (Python 3.7.4; https://www.python.org run on Windows 10) were used to compute angles and instantaneous velocities for all DeepLabCut-extracted markers, as well as the speed of the Bonsai-extracted centroid trajectories. All time series were applied a smooth filter (Hanning smoothing) of 10 frames (33 ms) before further processing.

#### Statistical analysis

All data are presented as mean ± standard error of mean (SEM). Data were analyzed using one-way analysis of variance (ANOVA), as appropriate. Bonferroni test was used for post-hoc analyses of significant ANOVAs to correct for multiple comparisons. Differences were considered significant at the level of p *<* 0.05. Statistical analysis was performed with Prism 9.0 (GraphPad, San Diego, CA).

## References

1. Grillner, S., and Manira, A. E. (2020). Current principles of motor control, with special reference to vertebrate locomotion. Physiological Reviews 100, 271–320. doi:10.1152/physrev.00012.2019.

2. Cullen, K. E. (2014). The neural encoding of self-generated and externally applied movement: implications for the perception of self-motion and spatial memory. Frontiers in Integrative Neuroscience 7. doi:10.3389/fnint.2013.00108.

3. Lewis, M. A., Fagan, W. F., Auger-Méthé, M., Frair, J., Fryxell, J. M., Gros, C., Gurarie, E., Healy, S. D., and Merkle, J. A. (2021). Learning and animal movement. Frontiers in Ecology and Evolution 9. doi:10.3389/fevo.2021.681704.

4. Schwaner, M. J., Hsieh, S. T., Braasch, I., Bradley, S., Campos, C. B., Collins, C. E., Donatelli, C. M., Fish, F. E., Fitch, O. E., Flammang, B. E., Jackson, B. E., Jusufi, A., Mekdara, P. J., Patel, A., Swalla, B. J., Vickaryous, M., and McGowan, C. P. (2021). Future tail tales: A forward-looking, integrative perspective on tail research. Integrative and Comparative Biology 61, 521–537. doi:10.1093/icb/icab082.

5. Fish, F. E., Rybczynski, N., Lauder, G. V., and Duff, C. M. (2021). The role of the tail or lack thereof in the evolution of tetrapod aquatic propulsion. Integrative and Comparative Biology 61, 398–413. doi:10.1093/icb/icab021.

6. Mincer, S. T., and Russo, G. A. (2020). Substrate use drives the macroevolution of mammalian tail length diversity. Proceedings. Biological sciences 287, 20192885. doi:10.1098/rspb.2019.2885.

7. Ehrlich, D. E. et al. (2017). Control of movement initiation underlies the development of balance. Current Biology 27, 334–344. doi:10.1016/j.cub.2016.12.037.

8. Shield, S., Jericevich, R., Patel, A., and Jusufi, A. (2021). Tails, flails, and sails: How appendages improve terrestrial maneuverability by improving stability. Integrative and Comparative Biology 61, 506–520. doi:10.1093/icb/icab108.

9. Lacava, S. A., Isilak, N., and Uusisaari, M. Y. (2024). The role of mice tails in response to external and self-generated balance perturbations on the roll plane. Journal of Experimental Biology jeb.247552. doi:10.1242/jeb.247552.

10. Murray, A. J., Croce, K., Belton, T., Akay, T., and Jessell, T. M. (2018). Balance control mediated by vestibular circuits directing limb extension or antagonist muscle co-activation. Cell Reports 22, 1325–1338. doi:10.1016/j.celrep.2018.01.009.

11. Goldberg, J. M., Wilson, V. J., Cullen, K. E., Angelaki, D. E., Broussard, D. M., Buttner-Ennever, J., Fukushima, K., and Minor, L. B. The Vestibular System: A Sixth Sense. Oxford University Press (2012). doi:10.1093/acprof:oso/9780195167085.001.0001.

12. Goldberg, J. M., and Cullen, K. E. (2011). Vestibular control of the head: possible functions of the vestibulocollic reflex. Experimental Brain Research 210, 331–345. doi:10.1007/s00221-011-2611-5.

13. Bath, A. P., Harris, N., and Yardley, M. P. (1998). The vestibulo-collic reflex. Clinical Otolaryngology Allied Sciences 23, 462–466. doi:10.1046/j.1365-2273.1998.00192.x.

14. Liang, H., Bácskai, T., Watson, C., et al. (2014). Projections from the lateral vestibular nucleus to the spinal cord in the mouse. Brain Structure and Function 219, 805–815. doi:10.1007/s00429-013-0536-4.

15. Highstein, S. M., and Holstein, G. R. The anatomy of the vestibular nuclei. In: Bü ttner-Ennever, J., ed. Progress in Brain Research vol. 151 (157–203). Elsevier. ISBN 9780444516961 (2006):(157–203). doi:10.1016/S0079-6123(05)51006-9.

16. Shinohara, H. (1999). The musculature of the mouse tail is characterized by metameric arrangements of bicipital muscles. Okajimas Folia Anatomica Japonica 76, 157–169. doi:10.2535/ofaj1936.76.4_157.

17. Biewener, A. A., and Blickhan, R. (1988). Kangaroo rat locomotion: Design for elastic energy storage or acceleration? Journal of Experimental Biology 140, 243–255. doi:10.1242/jeb.140.1.243.

18. Horak F, B. (2006). Postural orientation and equilibrium : what do we need to know about neural control of balance to prevent falls ? Age Aging (7–11). doi:10.1093/ageing/afl077.

19. Martín-Serra, A., Pérez-Ramos, A., Pastor, F. J., Velasco, D., and Figueirido, B. (2021). Phenotypic integration in the carnivoran backbone and the evolution of functional differentiation in metameric structures. Evolution Letters 5, 251–264. doi:10.1002/evl3.224.

20. Jones, K. E., Gonzalez, S., Angielczyk, K. D., and Pierce, S. E. (2020). Regionalization of the axial skeleton predates functional adaptation in the forerunners of mammals. Nature Ecology & Evolution 4, 470–478. doi:10.1038/s41559-020-1094-9.

21. Astley, H. C. (2020). The biomechanics of multi-articular muscle–tendon systems in snakes. Integrative and Comparative Biology 60, 140–155. doi:10.1093/icb/icaa012.

22. Zatsiorsky, V. M., and Prilutsky, B. I. Biomechanics of Skeletal Muscles. Human Kinetics (2012). doi:10.5040/9781492595298.

23. Mohan, R., Tosolini, A., and Morris, R. (2014). Targeting the motor end plates in the mouse hindlimb gives access to a greater number of spinal cord motor neurons: An approach to maximize retrograde transport. Neuroscience 274, 318–330. doi:10.1016/j.neuroscience.2014.05.045.

24. Henneman, E. (1957). Relation between size of neurons and their susceptibility to discharge. Science 126, 1345–1347. doi:10.1126/science.126.3287.1345.

25. D’Elia, K. P., and Dasen, J. S. (2018). Development, functional organization, and evolution of vertebrate axial motor circuits. Neural development 13, 1–12. doi:10.1186/s13064-018-0108-7.

26. Stifani, N. (2014). Motor neurons and the generation of spinal motor neuron diversity. Frontiers in Cellular Neuroscience 8. doi:10.3389/fncel.2014.00293.

27. Ohmori, Y., Watanabe, T., and Fujioka, T. (1984). Localization of the motoneurons innervating the hindlimb muscles in the spinal cord of the domestic fowl. Anatomia, Histologia, Embryologia 13, 141–155. doi:10.1111/j.1439-0264.1984.tb00582.x.

28. Ryan, M. J. et al. (1998). Topographic position of forelimb motoneuron pools is conserved in vertebrate evolution. Brain Behavior and Evolution 51, 90–99. doi:10.1159/000006401.

29. Toossi, A. et al. (2019). Functional organization of motor networks in the lumbosacral spinal cord of non-human primates. Scientific Reports 9, 13539. doi:10.1038/s41598-019-50041-3.

30. Levine, A. J., Lewallen, K. A., and Pfaff, S. L. (2012). Spatial organization of cortical and spinal neurons controlling motor behavior. Current Opinion in Neurobiology 22, 812–821. doi:10.1016/j.conb.2012.07.002.

31. Arber, S. (2017). Organization and function of neuronal circuits controlling movement. EMBO Molecular Medicine 9, 281–284. doi:10.15252/emmm.201607226.

32. Wilson, V. J., Boyle, R., Fukushima, K., Rose, P. K., Shinoda, Y., Sugiuchi, Y., and Uchino, Y. (1995). The vestibulocollic reflex. Journal of Vestibular Research: Equilibrium & Orientation 5, 147–170. doi:10.1016/0957-4271(94)00035-Z.

33. Takeoka, A., Vollenweider, I., Courtine, G., and Arber, S. (2014). Muscle spindle feedback directs locomotor recovery and circuit reorganization after spinal cord injury. Cell 159, 1626– 1639. doi:10.1016/j.cell.2014.11.019.

34. Lee, S., and Kaylie, D. M. Balance (anatomy: Vestibular nerve). In: Kountakis, S. E., ed. Encyclopedia of Otolaryngology, Head and Neck Surgery (577). Springer, Berlin, Heidelberg (2013): ( 577). doi:10.1007/978-3-642-23499-6_577.

35. Liang, H., Bácskai, T., Watson, C., and Paxinos, G. (2014). Projections from the lateral vestibular nucleus to the spinal cord in the mouse. Brain Structure and Function 219, 805– 815. doi:10.1007/s00429-013-0536-4.

36. Montardy, Q., Wei, M., Liu, X., Yi, T., Zhou, Z., Lai, J., Zhao, B., Besnard, S., Tighilet, B., Chabbert, C., and Wang, L. (2021). Selective optogenetic stimulation of glutamatergic, but not gabaergic, vestibular nuclei neurons induces immediate and reversible postural imbalance in mice. Progress in Neurobiology 204, 102085. doi:10.1016/j.pneurobio.2021.102085.

37. Henry, S. M., Fung, J., and Horak, F. B. (2001). Effect of stance width on multidirectional postural responses. Journal of Neurophysiology 85, 559–570. doi:10.1152/jn.2001.85.2.559.

38. Hong, S. L., Manor, B., and Li, L. (2007). Stance and sensory feedback influence on postural dynamics. Neuroscience Letters 423, 104–108. doi:10.1016/j.neulet.2007.06.043.

39. Goodworth, A. D., Mellodge, P., and Peterka, R. J. (2014). Stance width changes how sensory feedback is used for multisegmental balance control. Journal of Neurophysiology 112, 525–542. doi:10.1152/jn.00490.2013.

40. Sengupta, M., and Bagnall, M. W. (2023). Spinal interneurons: Diversity and connectivity in motor control. Annual Review of Neuroscience 46, 79–99. doi:10.1146/annurev-neuro-083122-025325.

41. Barmack, N. H. (2003). Central vestibular system: vestibular nuclei and posterior cerebellum. Brain research bulletin 60, 511–541. doi:10.1016/s0361-9230(03)00055-8.

42. Zhang, Y., Luo, Y., Sasamura, K., and Sugihara, I. (2021). Single axonal morphology reveals high heterogeneity in spinocerebellar axons originating from the lumbar spinal cord in the mouse. Journal of Comparative Neurology 529, 3893–3921. doi:10.1002/cne.25223.

43. Kodama, T., Guerrero, S., Shin, M., Moghadam, S., Faulstich, M., and du Lac, S. (2012). Neuronal classification and marker gene identification via single-cell expression profiling of brainstem vestibular neurons subserving cerebellar learning. Journal of Neuroscience 32, 7819–7831. doi:10.1523/JNEUROSCI.0543-12.2012.

44. Machado, A. S., Marques, H. G., Duarte, D. F., Darmohray, D. M., and Carey, M. R. (2020). Shared and specific signatures of locomotor ataxia in mutant mice. Elife 9, e55356. doi:10.7554/eLife.55356.

45. Nieuwenhuys, R. (1977). The brain of the lamprey in a comparative perspective. Annals of the New York Academy of Sciences 299, 97–145. doi:10.1111/j.1749-6632.1977.tb41902.x.

46. Amemiya, F., Kishida, R., Goris, R. C., Onishi, H., and Kusunoki, T. (1985). Primary vestibular projections in the hagfish, eptatretus burgeri. Brain research 337, 73–79. doi:10.1016/0006-8993(85)91610-5.

47. Fritzsch, B. (1998). Evolution of the vestibulo-ocular system. Otolaryngology–Head and Neck Surgery 119, 182–192. doi:10.1016/S0194-5998(98)70053-1.

48. Kubo, T., and Benton, M. J. (2009). Tetrapod postural shift estimated from permian and triassic trackways. Palaeontology 52, 1029–1037. doi:10.1111/j.1475-4983.2009.00897.x.

49. Harrison, M., O’Brien, A., Adams, L., Cowin, G., Ruitenberg, M. J., Sengul, G., and Watson, C. (2013). Vertebral landmarks for the identification of spinal cord segments in the mouse. NeuroImage 68, 22–29. doi:10.1016/j.neuroimage.2012.11.048.

50. Lö sel, P. D., et al. (2020). Introducing biomedisa as an open-source online platform for biomedical image segmentation. Nature Communications 11, 5577. doi:10.1038/s41467-020-19303-w.

51. Limaye, A. Drishti: A volume exploration and presentation tool. In: Proceedings of SPIE 8506, Developments in X-ray Tomography VIII. International Society for Optics and Photonics (2012):doi:10.1117/12.935640.

52. Jackman, S. L., Chen, C. H., Chettih, S. N., Neufeld, S. Q., Drew, I. R., Agba, C. K., Flaquer, I., Stefano, A. N., Kennedy, T. J., Belinsky, J. E., Roberston, K., Beron, C. C., Sabatini, B. L., Harvey, C. D., and Regehr, W. G. (2018). Silk fibroin films facilitate single-step targeted expression of optogenetic proteins. Cell Reports 22, 3351–3361. doi:10.1016/j.celrep.2018.02.081.

53. Fiederling, F., Hammond, L. A., Ng, D., Mason, C., and Dodd, J. (2021). Spineracks and spinalj for efficient analysis of neurons in a 3d reference atlas of the mouse spinal cord. STAR protocols 2, 100897. doi:10.1016/j.xpro.2021.100897.

54. Chiaruttini, N., Castoldi, C., Requie, L. M., Camarena-Delgado, C., dal Bianco, B., Gräff, J., Seitz, A., and Silva, B. A. (2024). Abba, a novel tool for whole-brain mapping, reveals brain-wide differences in immediate early genes induction following learning. bioRxiv. doi:10.1101/2024.09.06.611625.

55. Bankhead, P., Loughrey, M. B., Fernández, J. A., Dombrowski, Y., McArt, D. G., Dunne, P. D., McQuaid, S., Gray, R. T., Murray, L. J., Coleman, H. G., James, J. A., Salto-Tellez, M., and Hamilton, P. W. (2017). Qupath: Open source software for digital pathology image analysis. Scientific reports 7, 16878. doi:10.1038/s41598-017-17204-5.

56. Tyson, A. L., Vélez-Fort, M., Rousseau, C. V., Cossell, L., Tsitoura, C., Lenzi, S. C., Obenhaus, H. A., Claudi, F., Branco, T., and Margrie, T. W. (2022). Accurate determination of marker location within whole-brain microscopy images. Scientific Reports 12, 867. doi:10.1038/s41598-021-04676-9.

57. Stujenske, J. M., Spellman, T., and Gordon, J. A. (2015). Modeling the spatiotemporal dynamics of light and heat propagation for in vivo optogenetics. Cell Reports 12, 525–534. doi:10.1016/j.celrep.2015.06.036.

58. Bouvier, J., Caggiano, V., Leiras, R., Caldeira, V., Bellardita, C., Balueva, K., and Kiehn, O. (2015). Descending command neurons in the brainstem that halt locomotion. Cell 163, 1191–1203. doi:10.1016/j.cell.2015.10.074.

59. Mathis, A., Mamidanna, P., Cury, K. M., Abe, T., Murthy, V. N., Mathis, M. W., and Bethge, M. (2018). Deeplabcut: markerless pose estimation of user-defined body parts with deep learning. Nature Neuroscience. doi:10.1038/s41593-018-0209-y.

60. Lopes, G., Bonacchi, N., Frazão, J., Neto, J. P., Atallah, B. V., Soares, S., Moreira, L., Matias, S., Itskov, P. M., Correia, P. A., Medina, R. E., Calcaterra, L., Dreosti, E., Paton, J. J., and Kampff, A. R. (2015). Bonsai: an event-based framework for processing and controlling data streams. Frontiers in Neuroinformatics 9, 7. doi:10.3389/fninf.2015.00007.

