## Supplementary material for "A vestibulospinal pathway for context-dependent motor control of the mouse tail"

### SUPPLEMENTARY TABLES



Table 1: Anatomical measurements

| Parameter | mean±SEM | n | p-value (t-test) |
|---|---|---|---|
| **Soma size (MNs)** | | | p<0.0001, t-test |
| ext-MNs | $515.3 \pm 13.02$ | $n = 257$ | |
| int-MNs | $396.9 \pm 10.69$ | $n = 282$ | ext-MNs vs. int-MNs p<0.0001 |
| **Soma size (VC)** | | | p<0.0001, t-test |
| Nucleus X | $186.2 \pm 10.29$ | $n = 47$ | |
| Vestibular Nuclei | $670.9 \pm 17.54$ | $n = 171$ | Nucleus X vs. Vestibular Nuclei p<0.0001 |

Statistical analysis was performed using an unpaired t-test. $n$ refers to the number of data points.

Table 2: Vestibular Nucleus optogenetic stimulation

| Parameter | mean±SEM | n | p-value (t-test) |
|---|---|---|---|
| **Base of Support (45mm)** | | | p<0.0001, one-way ANOVA |
| $BoS_{L\text{-}BaseL}$ | $-11.18 \pm 4.92$ | $n = 7$ | |
| $BoS_{L\text{-}Opto}$ | $94.56 \pm 27.03$ | $n = 7$ | $BoS_{L\text{-}BaseL}$ VS $BoS_{L\text{-}Opto}$ p<0.0001 |
| $BoS_{R\text{-}BaseL}$ | $-5.22 \pm 1.44$ | $n = 7$ | |
| $BoS_{R\text{-}Opto}$ | $134.0 \pm 27.69$ | $n = 7$ | $BoS_{R\text{-}BaseL}$ VS $BoS_{R\text{-}Opto}$ p<0.0001 |
| **Base of Support (4mm)** | | | p<0.0001, one-way ANOVA |
| $BoS_{L\text{-}BaseL}$ | $6.048 \pm 4.42$ | $n = 7$ | |
| $BoS_{L\text{-}Opto}$ | $26.56 \pm 19.34$ | $n = 7$ | $BoS_{L\text{-}BaseL}$ VS $BoS_{L\text{-}Opto}$ p<0.0001 |
| $BoS_{R\text{-}BaseL}$ | $-1.115 \pm 4.59$ | $n = 7$ | |
| $BoS_{R\text{-}Opto}$ | $27.09 \pm 13.78$ | $n = 7$ | $BoS_{R\text{-}BaseL}$ VS $BoS_{R\text{-}Opto}$ p<0.0001 |
| **Tail rotation (%)** | | | p<0.01, one-way ANOVA |
| Tail $Rot_{L\text{-}Flat}$ | $12.70 \pm 5.77$ | $n = 7$ | |
| Tail $Rot_{L\text{-}Ridge}$ | $45.96 \pm 16.68$ | $n = 5$ | $BoS_{L\text{-}BaseL}$ VS $BoS_{L\text{-}Opto}$ p<0.039 |
| Tail $Rot_{R\text{-}Flat}$ | $1.43 \pm 1.43$ | $n = 5$ | |
| Tail $Rot_{R\text{-}Ridge}$ | $52.28 \pm 9.04$ | $n = 6$ | $BoS_{R\text{-}BaseL}$ VS $BoS_{R\text{-}Opto}$ p<0.0026 |

Statistical analysis was performed using paired t-test (comparison of two groups) or ANOVA (comparison of more than two groups). n refers to the number of datapoints. vs., versus.

Table 3: Sacral-Vestibulospinal optogenetic stimulation

| Parameter | mean±SEM | n | p-value (t-test) |
|---|---|---|---|
| **Tail Swing IL (%)** | | | |
| Tail swing $IL_{Opto}$ | $39.72 \pm 8.97$ | $n = 9$ | |
| Tail swing $IL_{Control}$ | $12.81 \pm 3.85$ | $n = 9$ | p=0.0092 |
| **Tail Swing CL (%)** | | | |
| Tail swing $CL_{Opto}$ | $26.95 \pm 7.27$ | $n = 11$ | |
| Tail swing $CL_{Control}$ | $16.69 \pm 5.40$ | $n = 11$ | p=0.071 |
| **Accuracy (no stimulation)** | | | |
| 10-degree | $0.66 \pm 0.068$ | $n = 7$ | |
| 8-degree | $0.54 \pm 0.098$ | $n = 7$ | p=0.14 |
| 5-degree | $0.19 \pm 0.044$ | $n = 7$ | p=0.016 |
| **Tail Swing IL/Stim Congruent (%)** | | | |
| Tail swing $IL_{Opto}$ | $50.14 \pm 8.34$ | $n = 9$ | |
| Tail swing $IL_{Control}$ | $23.14 \pm 6.53$ | $n = 9$ | p=0.0017 |
| **Tail Swing CL/Stim Congruent (%)** | | | |
| Tail swing $CL_{Opto}$ | $9.36 \pm 8.34$ | $n = 7$ | |
| Tail swing $CL_{Control}$ | $22.97 \pm 7.23$ | $n = 7$ | p=0.023 |
| **Tail Swing IL/Stim Non-Congruent (%)** | | | |
| Tail swing $IL_{Opto}$ | $32.45 \pm 6.09$ | $n = 7$ | |
| Tail swing $IL_{Control}$ | $22.68 \pm 6.69$ | $n = 7$ | p=0.0017 |
| **Tail Swing CL/Stim Non-Congruent (%)** | | | |
| Tail swing $CL_{Opto}$ | $23.82 \pm 6.58$ | $n = 10$ | |
| Tail swing $CL_{Control}$ | $19.99 \pm 4.27$ | $n = 10$ | p=0.56 |

Statistical analysis was performed using paired t-test (comparison of two groups) or ANOVA (comparison of more than two groups). n refers to the number of datapoints. vs., versus.

### SUPPLEMENTARY FIGURES

Figure 1: **Tail-motoneurons targeting the lateral extrinsic muscle and AAV9 tropism.**

**A1:** Schematic depiction of the approach for locating lateral tail-motoneurons. Red and green indicate extrinsic (lateral and ventral respectively) muscles; orange represents distal (intrinsic) muscles.
**A2:** Example results of retrograde labeling with injection to lateral extrinsic tail muscles, shown as standard deviation projection images (20x).
**B:** Example results of two retrogradely labeled motoneurons, one labeled with CTB (dotted green) and the other with AAV9 virus (solid green). Based on soma size, they were identified as putative alpha and gamma motoneurons, respectively, following criteria in Friese et al. (2009).
**C:** Histogram of lateral tail motoneurons soma size labeled with CTB (light green, $n = 120$, $N = 4$ mice) or AAV (dark green, $n = 31$, $N = 4$ mice).
**D1-D2:** Density maps of lateral tail-motoneuron locations from 4 mice. The green line represents the lateral tail motoneuron distribution in sacral 3 (D1) and sacral 4 (D2), while the black line shows the distribution of the entire extrinsic tail motoneuron pool.

Figure 2: **Transynaptic tracing shows projections from the lumbar spinal cord and the lateral vestibular nucleus to tail motoneurons.**

**A:** Tail motoneurons labeled via viral injection into extrinsic tail muscles, following standard protocol. Scale bar = 200 µm.
**B:** Representative sections through lumbar spinal cord, showing projection neurons to tail-MNs following rabies virus labeling. Scale bar = 500 µm.
**C:** Representative sections through lateral vestibular nucleus, with labeled vestibulospinal neurons contacting tail motoneurons. Subnuclei boundaries indicated. Scale bar = 500 µm.

Figure 3: **VN optogenetic stimulation effect on the base of support in the ridge and forward speed**

**A:** Comparison of the BoS-change responses to left and right stimulation while mice cross the ridge (N=7).
**B:** Effect of optogenetic activation on forward speed in the ridge (green), and flat surface (blue).

Figure 4: **sacVS optogenetic stimulation effect on tail and BoS.**

**A1-3:** Example of single trial from one mouse of the tail response to optogenetic stimulation (blue shaded area) in response to VN stimulation (A1), sacVS stimulation (A2), or no stimulation (sacVS control, A3). Grey traces are trials with no tail swing and pink traces with tail swing.
**B-C:** Effect of optogenetic activation on the base of support in the ridge (green), and flat surface (orange) without stimulation (B), and with optogenetic stimulation (C).

Figure 5: **Effect of tilt decrease in accuracy measure and sacVS effect on tail kinematic.**

**A:** Decrease in accuracy is shown for trials where roll-tilt applied was smaller in amplitude (10, 8 and 5 degree). One-way ANOVA, N=7. * $p<0.5$.
**B1-2:** Distribution of tail swing reaction time (B1, bin-width = 20), and maximum velocity (B2, bin-width = 3) across trials with optogenetic (pink) or control (grey).

### Video captions

**Supplementary video 1.** Tail base muscle reconstruction using microCT scan acquisition. Coccygeal vertebrae are shown from Co2 to Co10. Extrinsic muscles are shown in the coronal section with blue (dorsal), red (lateral) and green (ventral). The intrinsic muscles are shown in orange, purple, and light green (dorsal, lateral and ventral respectively). The lateral tendon is depicted in yellow.
**Supplementary video 2.** Optogenetic stimulation of right VN while mouse crossed a flat surface. Following a trial with activation of right VN in the same mouse on the ridge. The onset of optogenetic stimulation is indicated by LED turning on on the bottom right corner.
**Supplementary video 3.** Example of ipsilateral trials showing the tail response to roll tilt of different intensities (10 and 5 degrees).
**Supplementary video 4.** Optogenetic stimulation of sacVS while mouse crossed the ridge (slow motion). Following a control trial where the same mouse is crossing the ridge without the optogenetic stimulation (time-window to the optogenetic trial is indicated by the red LED activation in the bottom right corner).
